# Conserved allosteric inhibition mechanism in SLC1 transporters

**DOI:** 10.1101/2022.09.21.508810

**Authors:** Yang Dong, Jiali Wang, Rachel-Ann Garibsingh, Keino Hutchinson, Yueyue Shi, Gilad Eisenberg, Xiaozhen Yu, Avner Schlessinger, Christof Grewer

## Abstract

Excitatory Amino Acid Transporter 1 (EAAT1) is a plasma-membrane glutamate transporter belonging to the SLC1 family of solute carriers. It plays a key role in neurotransmitter transport and contributes to the regulation of the extracellular glutamate concentration in the mammalian brain. The structure of EAAT1 was determined using cryo-EM, in complex with UCPH-101, a highly potent and non-competitive inhibitor of EAAT1. Alanine Serine Cysteine Transporter 2 (ASCT2) is a neutral amino acid transporter, which regulates pools of amino acids such as glutamine, serine and alanine between intracellular and extracellular compartments in a Na^+^ dependent manner. ASCT2 also belongs to the SLC1 family and shares 58% sequence similarity with EAAT1. However, allosteric modulation of ASCT2 via non-competitive inhibitors is unknown. Here we explore the UCPH-101 inhibitory mechanisms of EAAT1 and ASCT2 by using rapid kinetic experiments. Our results show that UCPH-101 slows substrate translocation rather than substrate or Na^+^ binding, confirming a non-competitive inhibitory mechanism, but only partially inhibits wild-type ASCT2 with relatively low affinity. Guided by computational modeling using ligand docking and molecular dynamics (MD) simulations, we selected two residues involved in UCPH-101/EAAT1 interaction, which were mutated in ASCT2 (F136Y, I237M, F136Y/I237M) in the corresponding positions. We show that in the F136Y/I237M double mutant transporter, 100% of the inhibitory effect of UCPH-101 on anion current could be restored, and the apparent affinity was increased (*K*_i_ = 9.3 μM), much closer to the EAAT1 value of 0.6 μM. Finally, we identify a novel non-competitive ASCT2 inhibitor, identified through virtual screening and experimental testing against the allosteric site, further supporting its localization. Together, these data indicate that the mechanism of allosteric modulation is conserved between EAAT1 and ASCT2. Due to the difference in binding site residues between ASCT2 and EAAT1, these results raise the possibility that more potent, and potentially selective inhibitors can be designed that target the ASCT2 allosteric binding site.

## Introduction

Excitatory amino acid transporters (EAATs) from the solute carrier 1 (SLC1) family are an important class of membrane proteins, which are strongly expressed in the mammalian central nervous system (CNS). EAATs are responsible for the transport of glutamate across neuronal and astrocytic membranes in the CNS. Glutamate transporters are secondary active transporters, which utilize energy from the sodium concentration gradient between the extra- and intracellular sides of the membrane to take up glutamate into cells against a large concentration gradient (1,2). This glutamate uptake is an important process, preventing glutamate concentrations from reaching neurotoxic levels in the extracellular space (2,3).

The SLC1 family contains five glutamate transporters (EAATS1-5), as well as two neutral amino acid transporters, the alanine serine cysteine transporters ASCT1 and 2 (4-6). ASCT2 has been implicated in rapidly-growing cancer cells as a supply mechanism for glutamine, and inhibition of the transporter was shown to reduce cell proliferation and tumor size (7-10). Therefore, ASCT2 is emerging as a promising drug target.

Structurally, SLC1 family members are assembled from three identical subunits (11-13), each contain 8 transmembrane domains (TM1 to TM8). TM1 to TM6 form the trimerization domain, and TM7, TM8 together with hairpin loops HP1 and HP2 form the transport domain, which is based on an inverted repeat structure (14,15). Glutamate transporter inhibitors were widely studied for pharmacological purposes. Several types of competitive inhibitors were developed based on an aspartate scaffold, for example TBOA (DL-*threo*-beta-benzyloxyaspartate) (16) and TFB-TBOA ((3*S*)-3-[[3-[[4-(Trifluoromethyl)benzoyl]amino]phenyl]methoxy]-L-aspartic acid) (17). In addition, non-competitive inhibitors were identified such as UCPH-101 and 102 (18,19). Characterization with electrophysiological methods suggested UCPH-101 was specific for EAAT1 rather than EAATs 2-5. The inhibition of EAAT1 by UCPH-101 was not affected by glutamate or competitive inhibitors, such as TBOA (16). Subsequently, the binding site of UCPH-101 was identified, first by site-directed mutagenesis, and then in a UCPH-101-bound crystal structure (11). UCPH-101 was found to bind to a hydrophobic region between the scaffold and transport domains (11).

In addition to UCPH-101, other allosteric modulators, GT949 and derivatives, were first introduced by the Falucci et al. (20,21). GT949 is a positive allosteric modulator in EAAT2. However, it operates as an inhibitor in other EAAT subtypes. After incubation of GT949, glutamate uptake activity and *v*_max_ increase in EAAT2, without altering the substrate affinity. The report also suggested that the allosteric binding site is located between the transport and the trimerization domains (20,21).

For ASCTs, inhibitor pharmacology is less well studied. Currently, a number of competitive inhibitors have been identified, on the basis of structural similarity with EAAT inhibitors, as well as docking to homology models (22,23). While these newly discovered substrate binding site inhibitors show improved potencies and selectivities (24), because of their high similarity to amino-acid substrates they (i) likely have poor bioavailability and (ii) need to compete with the amino acid rich media; thus, they are unlikely to be useful as future anti-cancer drugs via inhibition of ASCT2-mediated transport. It is therefore critical to identify specific, allosteric, non-amino-acid like inhibitors for ASCT2. EAAT1 and ASCT2 share 57.6% sequence similarity and 38.5% sequence identity (Fig. S1); however, it is not known if the allosteric binding site of EAAT1 is also found in ASCTs, and whether small molecules can modulate ASCT2 non-competitively via this potential site. Structural analysis of the allosteric sites of EAAT1 and ASCT2 suggests that two non-conserved residues in EAAT1 (e.g., F136, and I237) significantly contribute to the UCPH-101 interaction, while other binding site residues that are less critical for binding are conserved between EAAT1 and ASCT2 (Fig. S1).

In this report, we first compare and detail the effects of UCPH-101 on the SLC1 family members EAAT1 and ASCT2, including analysis of rapid chemical kinetic experiments. While UCPH-101 was found to be a partial and low affinity inhibitor of wild-type ASCT2, engineering of a double mutant ASCT2 transporter based on computational docking analysis on the basis of the EAAT1 binding site, resulted in a largely recovered UCPH-101 effect. In addition, we describe the identification of a non-competitive inhibitor that is not related to UCPH-101 from a virtual screen. Finally, we discuss the importance of our results to the understanding of allosteric inhibition in SLC1 transporters as well as their relevance to the identification of future, non-competitive modulators for ASCTs, including anti-cancer drugs.

## Materials and Methods

### Cell Culture and Transfection

HEK293 cells (American Type Culture Collection No. CRL 1573) were cultured as described previously(25). Cell cultures were transiently transfected with wild-type EAAT1 or wild-type/mutant rASCT2 cDNAs, inserted into a modified pBK-CMV-expression plasmid. Transfection were performed according Jetprime transfection reagent and protocol (Polyplus). The cells were transfected after 24-36 hours then used for electrophysiological analysis.

### Electrophysiology

Currents associated with ASCT2 and EAAT1 were measured in the whole-cell current recording configuration. Whole-cell currents were recorded with an EPC7 patch-clamp Amplifier (ALA Scientific, Westbury, NY) under voltage-clamp conditions. The resistance of the recording electrode was 3-6 MΩ. Series resistance was not compensated because of the small whole-cell currents carried by EAAT1, ASCTs. The composition of the solutions for measuring amino acid exchange currents in the anion conducting mode was: 140 mM NaMes (Mes = methanesulfonate), 2 mM MgGluconate_2_, 2 mM CaMes_2_, 10 mM 4-(2-Hydroxyethyl)piperazine-1-ethanesulfonic acid (HEPES), with additional amino acid substrate (Glu or Ser) pH 7.3 (extracellular), and 130 mM NaSCN (SCN = thiocyanate, for EAAT1 NaSCN was replaced with KSCN), 2 mM MgGluconate_2_, 5 mM Ethylene glycol-bis(2-aminoethylether)-N,N,N’,N’-tetraacetic acid (EGTA), 10 mM HEPES, pH 7.3 (intracellular), as published previously (26). For the measurement of the transport component of the current, intracellular SCN^-^ was replaced with the Mes^-^ anion.

### Voltage-Jump Experiments

Voltage jumps (−100 to +60 mV) were applied to perturb the translocation equilibrium and to determine the voltage dependence of the anion conductance (27). To determine EAAT1/ ASCT-specific currents, external solution contained 140 mM NaMes in the presence of varying concentrations of amino acid substrate. The internal solution contained 130 mM NaSCN (KSCN for EAAT1) in the presence of 10 mM amino acid substrate. Competitive blocker TBOA(16,17), (R)-gamma-(4-biphenylmethyl)-L-proline (22) or UCPH-101 was used in control voltage jump experiments, yielding the unspecific current component, which was subtracted from the total current. Capacitive transient compensation and series resistance compensation of up to 80% was employed using the EPC-7 amplifier. Non-specific transient currents were subtracted in Clampfit software (Molecular Devices).

### Rapid Solution Exchange

Fast solution exchanges were performed using the SF-77B (Warner Instruments, LLC, MA, USA) piezo-based solution exchanger, allowing a time resolution in the 10 ms range. Amino acid substrate was applied through a theta capillary glass tubing (TG200-4, OD = 2.00 mm, ID = 1.40 mm. Warner Instruments, LLC, MA, USA), with the tip of the theta tubing pulled to a diameter of 350 μm and positioned at 0.5 mm to the cell (25,28). For paired-pulse experiments, currents were recorded with 10/20ms interval time after removal of amino acid.

### Laser photolysis

Laser-pulse photolysis experiments were performed as described in detail previously (29,30). In the reverse transport experiments, MNI-caged glutamate was introduced into the cell through the glass recording electrode. After whole-cell mode was established, a period of 5mins was allowed for diffusive equilibration with the cell interior. MNI-caged glutamate (31,32), at concentrations of 1 mM were applied to the cells and photolysis of the caged glutamate was initiated with a light flash (355nm, 8ns, Minilite II, Continuum). The light was coupled into a quartz fiber (diameter 365 μm) that was positioned in front of the cell in a distance of 300 μm. With maximum light intensities of 500–840 mJ/cm^2^ saturating glutamate concentrations could be released.

### Amino acid uptake

HEK293T cells were plated on collagen-coated 24-well plates (0.5 × 10^5^ cells/well) in Dulbecco’s modified Eagle’s medium containing 10% fetal bovine serum, penicillin (100 units/ml), 4.5 g/L glucose, sodium pyruvate, and glutamine (4 mM). 48 h post transfection with rASCT2 (see above), the cells were washed with uptake buffer (140 mM sodium methanesulfonate (NaMes), 2 mM magnesium methanesulfonate, 2 mM calcium gluconate, 30 mM HEPES, pH 7.4, 5 mM glucose) 2 times and preincubated in the same buffer for 10 min. Subsequently, buffer was replaced with fresh buffer containing unlabeled L-serine and 0.1 Ci of [^14^C] L-serine (Moravek Biochemicals; total concentration 100 μCi/mL). After 5 min of incubation at room temperature, uptake was terminated by washing twice with 0.1 ml of uptake buffer on ice (after 5 min, uptake was in the linear range, as determined by quantifying the time dependence of uptake for times up to 5 min). The cells were then solubilized in 0.2 mL of 1% SDS, and radioactivity was measured by scintillation counting in 3 mL of scintillation fluid. Unspecific L-serine uptake was determined using the ASCT2 competitive inhibitor L-cis-BPE (24).

### Molecular dynamics (MD) simulations

The model system for MD simulations was generated with VMD software (33) using an EAAT1 (5LLM) and ASCT2 (7BCS) structure. The final EAAT1/ASCT2 model, was inserted in a pre-equilibrated POPC lipid bilayer with the dimensions about 130 × 130 × 90 Å. TIP3P water was added to generate a box measuring about 100 Å in the z-direction. NaCl were added at a total concentration of 0.15 M and the system was neutralized. The total numbers of atoms in the EAAT1 system were 168926, 193804 in ASCT2. Simulations were run using the CHARMM36 force field. NAMD (34) simulations were performed using 2000 steps of minimization, followed by 10 ns equilibration runs under constant pressure conditions (NPT), and then for 100ns (35). The RMSD increased from 1.5 Å soon after simulation start to ∼3 Å after 2 ns of equilibration, after which it was in steady state. The cutoff for local electrostatic and van der Waals interactions was set to 12 Å. For long-range electrostatic interactions, we used the particle-mesh Ewald method implemented in NAMD. Bonds to hydrogen atoms and TIP3P water were kept rigid using SHAKE. The time steps of the simulations were 2 fs.

### Data Analysis

The data analysis was performed in Microsoft Excel and Microcal Origin software. Error bars are shown as mean ± standard deviation, collected from recordings of 6 to 10 cells, for statistical analysis. To determine the recovery rate, non-linear curve fitting was used with a Michaelis-Menten like equation, *I* = *I*_max_ * [inhibitor] / (*K*_i_ + [inhibitor]), where *I*_max_ is the current at saturating substrate concentration. Transient signals of piezo-based solution-exchange results were analyzed in Clampfit software (Axon Instruments) by fitting with a sum of two exponential components. *I* = *I*_1_·exp(-t/τ_rise_) + *I*_2_·exp(-t/τ_decay_). Here, *I* is the current amplitude, τ the time constant, and t the time.

### Molecular docking

All docking calculations were conducted with the Schrödinger package (36). ASCT2 (PDB ID: 7BCS) was prepared for docking using the Maestro Protein Preparation Wizard under default parameters. Glide v2020-2 was used to perform docking. The allosteric binding site was defined using the Maestro Receptor Grid Generation panel and the coordinates of the site were marked with UCPH-101, a reference ligand extracted from a superposition of an EAAT1 structure solved with bound UCPH-101 (PDB ID:5LLM or 5MJU generated the same result). We performed both rigid and flexible docking with and without hydrogen bond constraints at F377 and positional constraints in the non-mutated ASCT2 structure. However, docking poses were unrealistic and exhibited poor scores, and we thus, concluded they are unusable for further study.

### Docking UCPH-101 to mutated ASCT2

We remodeled the sidechains of two residues in ASCT2 (i.e. F136Y and I273M) on a fixed backbone to mimic the UCPH-101 binding site in EAAT1, using SCWRL4 (37). Grid preparation utilized positional constraints as highlighted in Fig. 1A and excluded volume at F192 with a radius of 2.5 Å. The pose that had the best docking score was used to carry out the MD simulations.

**Fig. 1:**
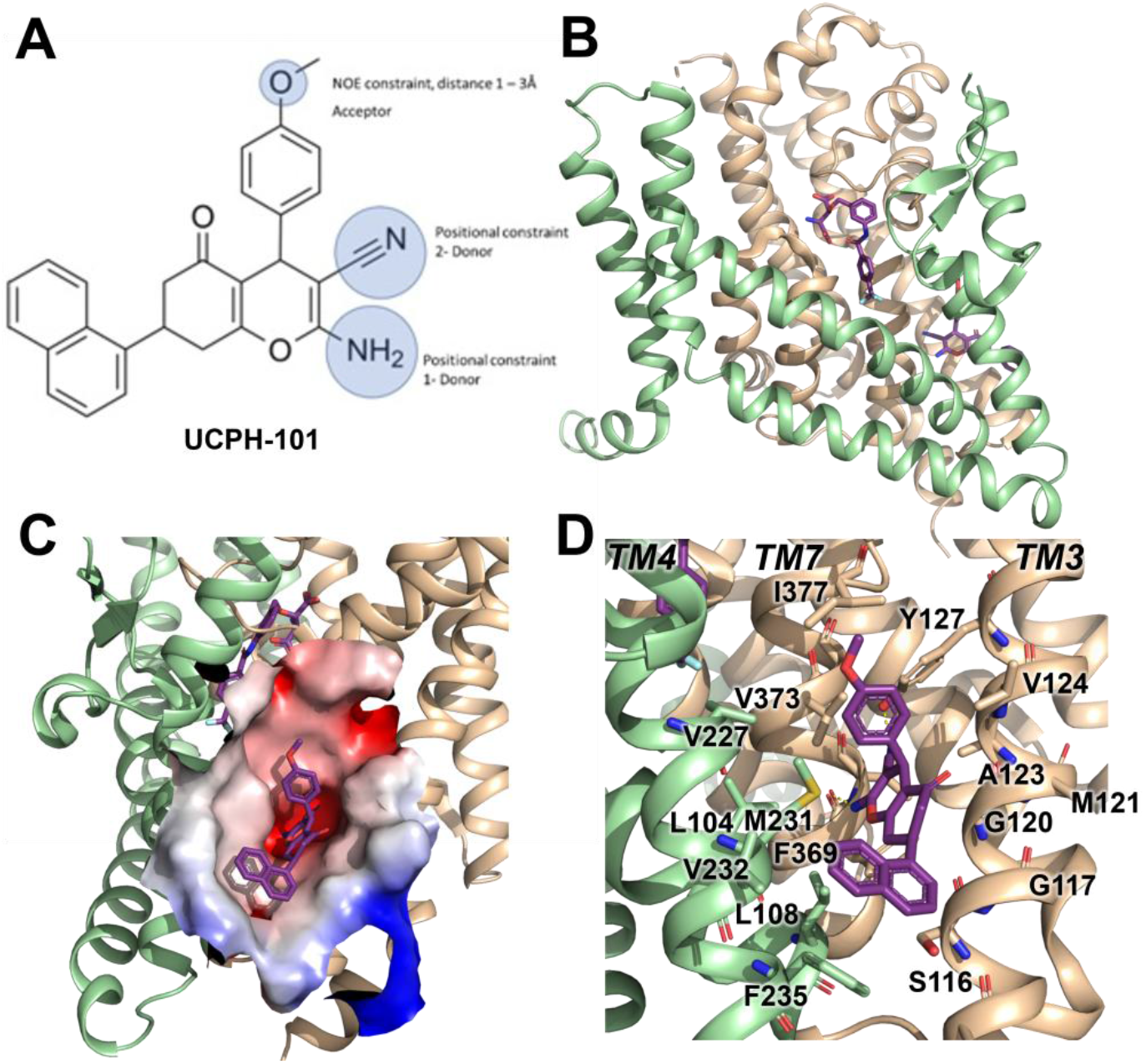
Structure of the EAAT1 UCPH-101-bound state. **(A)** UCPH-101 structure and substituents used for constraints in docking calculations. **(B)** EAAT1 structure (5LLM) in complex with TFB-TBOA in the substrate binding site and UCPH-101 in the allosteric site. The trimerization domain is highlighted in green, the transport domain in brown. **(C)** Illustration of the UCPH-101 binding site at the domain interface. The electrostatic surface of the binding site is highlighted. **(D)** EAAT1 transmembrane (TM) helices and amino acid residues in proximity of the UCPH-101 binding site.

### Virtual screening

The ZINC20 Lead-Like library (‘In Stock’; 3.8 million compounds) was virtually screened against the WT ASCT2 allosteric site using Glide, as described above. The 1,000 top-scoring compounds from the docking screen were further analyzed. In particular, because errors in docking can occur in large virtual screens, we inspected the docking poses of the top-ranking compounds to remove molecules with problematic pose or strained conformations, and prioritized them for experimental testing (38). We focused on molecules that interact with the conserved residues in the allosteric site (Fig. 2C). We purchased 11 molecules for experimental testing.

**Fig. 2:**
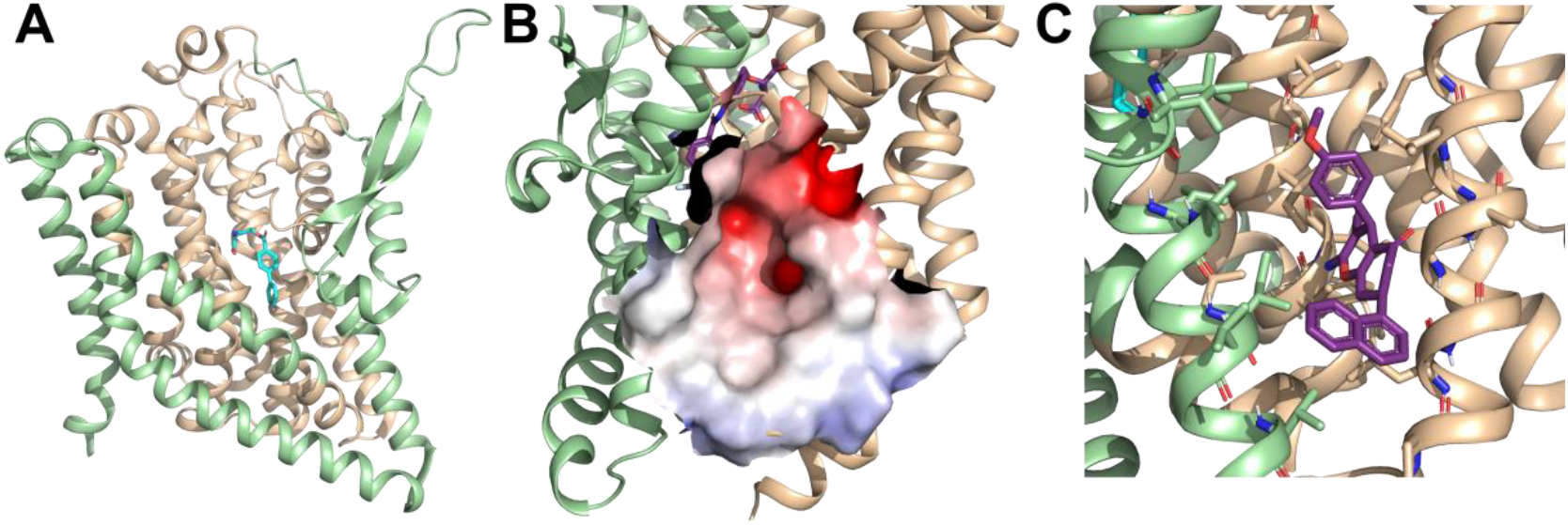
ASCT2 structures and predicted allosteric binding site. (**A)** The ASCT2 structure (PDB id 7BCS) in complex with the UCPH-101 coordinates derived from the EAAT1 structure in the allosteric site. The trimerization domain is highlighted in green, the transport domain in brown. **(B)** Illustration of the UCPH-101 binding site at the domain interface. The electrostatic surface of the binding site is highlighted. **(C)** Close-up view of the predicted ASCT2 UCPH-101 binding site.

## Results

UCPH-101 has been previously reported to be a potent non-competitive glutamate transporter inhibitor with high specificity for EAAT1 rather than the other SLC1 glutamate transporter subtypes. The EAAT1 structure shows UCPH-101 (Fig. 1A) deeply buried between the transport domain and the scaffold domain interface (Figs. 1B and C). The hydrophobic pocket is located between TM3, TM7, and TM4 (Fig. 1C, see also Fig. 4A, below). The residues that interact with UCPH-101 include G120 (TM3), V373 (TM7a), M231 (TM4c), Y127 (TM3), F369 (TM7) and F235 (TM4c), Fig. 1D. This binding pocket is at a distance of over 15 Å from the substrate (Fig. 1B) or cation binding sites, suggesting that UCPH-101 may not preclude substrate binding (11).

A comparison of the UCPH-101 binding pocket between EAAT1 and ASCT2 (Figs. 1B, 2A and C) reveals similarities and differences in the overall shape and physical chemical properties of the allosteric binding site of EAAT1 and the equivalent region in ASCT2 (Figs. 1C and 2B). Several residues are conserved between these proteins maintaining the overall shape of the site. For example, L104, V227, F369 in EAAT1 correspond to L112, V235, and F377 in ASCT2. Conversely, some residues are not fully conserved. Most notably, Y127 and M271 correspond to F136 and I237 in ASCT2, affecting the shape and polarity the binding site (Figs. S1 and 2B, C). We hypothesized that these differences as well as other changes will lead to differences in inhibitor specificity of the corresponding allosteric sites.

Indeed, molecular docking of UCPH-101 to the putative allosteric site of ASCT2 fails using multiple approaches including both rigid and flexible docking, as well as constraints on different interactions and residues (Methods). However, when F136 and I273 are remodeled to tyrosine and methionine respectively, UCPH-101 can be successfully docked into ASCT2 with a pose similar to that seen in the EAAT1 structures (see below). This result added further support to our hypothesis that Y127 and M271 are key residues impacting binding of UCPH-101 in EAAT1 and explains why inhibition is only partial in wild type ASCT2 (see below). Next, our goal was to experimentally test predictions from docking with respect to interaction of UCPH-101 with the SLC1 family member ASCT2, contrasting them with results from EAAT1. Therefore, we first describe mechanistic analysis of the UCPH-101 effect on EAAT1, which also revealed some novel mechanistic information.

### UCPH-101 is a non-competitive inhibitor of EAAT1

We applied whole-cell current recording using EAAT1/ASCT2-transfected HEK293 cells to study the UCPH-101 inhibition mechanism. Glutamate application (100 μM, K^+^ in the pipette and Na^+^ in the extracellular solution) induced large inwardly-directed anion currents (black) due to the outflow of internal SCN^-^ (Fig. 3A). For wild-type EAATs, glutamate-induced anion current was previously shown to be an indicator of transport activity (30,39). This glutamate-induced anion current was inhibited in a concentration-dependent manner after adding UCPH-101 to the extracellular solution. 2 μM UCPH (Fig. 3A red) resulted in over 50 % inhibition of anion currents. With up to 50 μM concentration of UCPH-101, the anion conductance was fully blocked (Fig. 3A; blue).

**Fig. 3:**
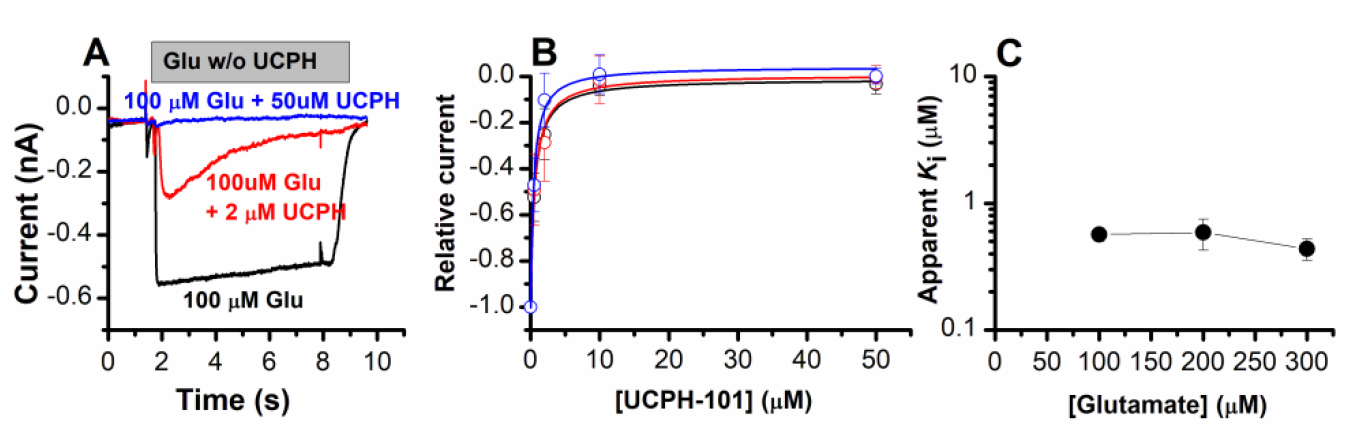
UCPH-101 is a high-affinity, non-competitive inhibitor of EAAT1 anion current. **(A)** Typical whole-cell current recordings in the absence (black) and presence of 2 μM (red) and 100 μM (blue) UCPH-101. [glutamate] was 100 μM. Experiments were performed using 140 mM NaMes in the extracellular buffer, and 130mM KSCN intracellularly (forward transport conditions). **(B)** Dose response curves to determine the apparent *K*_i_ for UCPH-101 at increasing glutamate concentration of 100 μM (black, n=16), 200 μM (red, n=19) and 300 μM (blue, n=13). **(C)** Glutamate concentration-dependence of the apparent *K*_i_ for UCPH-101 suggests non-competitive inhibition mechanism. All experiments were performed at 0 mV membrane potential.

To confirm the previously-proposed non-competitive inhibition mechanism of UCPH-101, we next determined the apparent inhibition constant (*K*_i_) for UCPH-101 with glutamate concentrations varying from 100 μM to 300 μM. Apparent inhibition constants (*K*_i_) were obtained by fitting the dose response curve of the current (*I*) with the following equation:

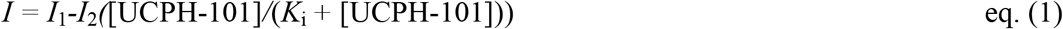

Here, *I*_1_ and *I*_2_ are the current amplitudes in the absence of inhibitor, and at saturating UCPH-101 concentration. Based on the expected non-competitive binding pattern, the UCPH-101 inhibition effect should not be dependent on the substrate concentration. In agreement with this expectation, the apparent *K*_i_ was 0.57 ± 0.07 μM, 0.59 ± 0.16 μM, and 0.44 ± 0.08 μM, at 100 μM, 200 μM, 300 μM glutamate concentration, respectively (Fig. 3B). By plotting the concentration dependent apparent *K*_i_ values (Fig. 3C), increasing the [glutamate] did no significantly change the *K*_i_ for UCPH-101. In contrast, if UCPH-101 would show a competitive inhibition pattern, an increase of apparent *K*_i_ with the glutamate concentration would be expected, according to eq. (2).

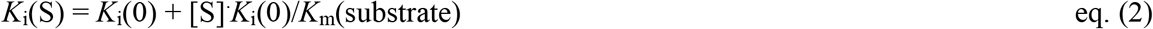

In eq. (2), *K*_i_(0) and *K*_i_(S) are the *K*i values (inhibition constants) in absence and presence of glutamate. *K*_m_(substrate) is the apparent Michaelis-Menten constant for glutamate, and [S] is the substrate concentration.

Next, we determined the voltage dependence of the UCPH-101 inhibitory effect. For this purpose, 140 mM Na^+^ with 1 mM glutamate, in the absence or presence of 50 μM UCPH were applied in the external solution to the transporter in the presence of intracellular SCN^-^. A voltage jumps from 0 mV to a final value ranging from +60 mV to −100 mV (voltage protocol shown in Fig. S2A, top panel) was applied to the cells. As expected, negative voltages resulted in larger anion currents, due to the increased driving force for SCN^-^ outflow. After incubation with a saturating concentration of UCPH-101, anion currents were fully inhibited at all membrane potentials (Figs. S2B and C).

### EAAT1 pre-steady-state kinetics in the presence of UCPH-101

To obtain a high time resolution for determining the effect of UCPH-101 on transport kinetics, we applied glutamate rapidly to EAAT1 using a piezo-based solution exchange device. This system has a 5-10 ms time resolution when applied to whole cells, and also allows the rapid removal of substrate, providing information on the recovery of current after glutamate removal. However, due to the millisecond time resolution, it will limit determination of very early intermediates in the transport cycle, which are present in a sub-millisecond time domain, but the time resolution is high enough for determining glutamate transporter turnover rate, and how it is affected by the presence of inhibitor. EAAT1 anion current responses to application of 1mM glutamate in two successive 20 ms pulses with variable spacing in between pulses (Fig. 4A) show rapid rise of the anion current after glutamate application within the time resolution afforded by solution exchange (5-10 ms). The rise of the current is followed by rapid decay of a transient anion current component to the steady state, as demonstrated previously (30). This phase of the current was previously assigned to steps associated with the glutamate translocation reaction (27). When glutamate was removed, the decay of the steady-state current was much slower, with a time constant of 32 ms (Fig. 4A), as had been reported previously for several glutamate transporter subtypes (30,40-42). After pre-incubation and in the continuous presence of UCPH-101, current amplitude was reduced in a dose-dependent manner, as expected (Fig. 4B, current was fully eliminated at 40 μM UCPH-101), but at concentrations close to the apparent *K*_i_, the decay of the current was also slowed (Fig. 4D, τ = 11 ± 1.3 ms, 16 ± 2.1ms, 20 ± 2.1ms, at 0 μM, 0.2 μM and 2 μM UCPH-101, respectively). In contrast to competitive inhibitors, UCPH-101 application in the absence of glutamate did not induce any currents, as expected for non-competitive inhibitors, which are not known to be able to block the glutamate transporter leak anion conductance (Fig. 4C).

**Fig. 4:**
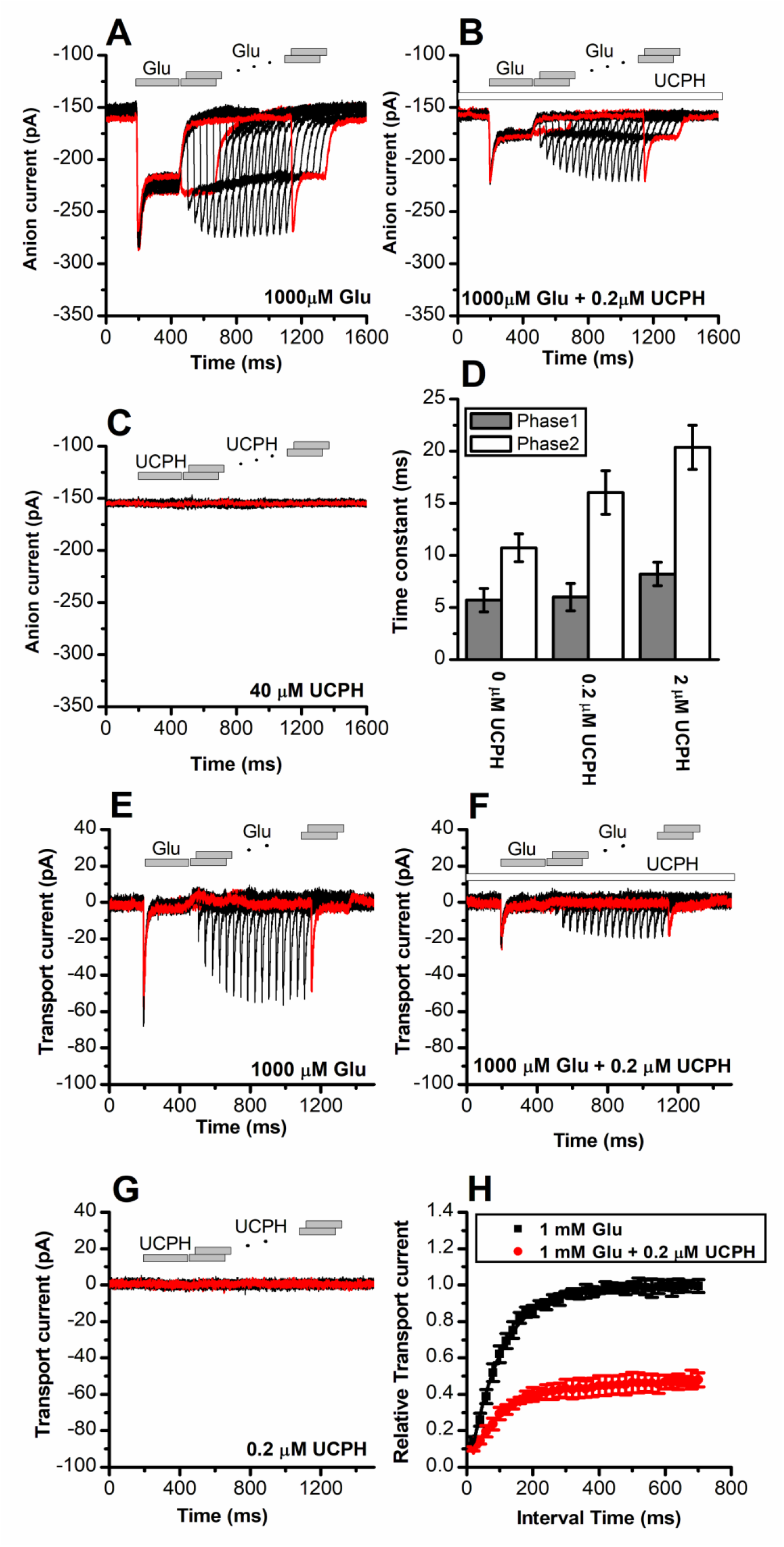
UCPH-101 has only minor effect on recovery kinetics of EAAT1 current after glutamate removal. **(A)** Anion currents in response to two pulses of rapid glutamate application (1 mM), with varying inter-pulse interval (pulse protocol shown at the top) under forward transport conditions. The intracellular solution contained 130 mM KSCN, the extracellular solution contained 140 mM NaMes. **(B)** Similar experiment as in (A), but in the presence of 0.2 μM UCPH-101 (pre-incubated for 5mins, see open bar for timing of solution exchange, top of the figure). **(C)** Application of UCPH-101 alone did not induce any currents. **(D)** Time constants for the fast and slow phase of the current recovery, for the two exponential components. **(E-G)** Experiments similar to (A-C), but for the transport component of the current (the permeant intracellular anion, SCN^-^, was replaced with the non-permeant Mes^-^ anion). **(H)** Recovery of the transient current in the presence and absence of 0.2 μM UCPH-101. The membrane potential was 0 mV in all experiments.

### UCPH-101 blocks steps associated with EAAT1 glutamate translocation

We next recorded glutamate-induced pre-steady-state currents under transport conditions, i.e. in the presence of non-permeable anion, using Mes^-^ to replace the permeable anion SCN^-^. As reported previously (29,43), transient inward currents were observed in response to glutamate concentration jumps, followed by a small steady-state component (Fig. 4E). The transient current was previously assigned to inward charge movement from a combination of Na^+^ binding and glutamate translocation (29,43). Upon removal of glutamate, the current rapidly decays to the baseline with little positive overshoot (in contrast to ASCT2, see below). Recovery of the transient current upon glutamate application in a second pulse is relatively slow, with a time constant of 95±4.2 ms, most likely reflecting the turnover time of the transporter, which has to recycle its binding sites to outward-facing, after passing through a whole transport cycle. In the presence of UCPH-101, both pre-steady state and steady-state currents are reduced in a dose-dependent manner (Figs. 3E,F). Here, EAAT1 was pre-incubated with UCPH-101 for 5 min. At 0.2 μM UCPH-101 recovery of the transient current after glutamate removal occurred with a time constant of 115±11 ms, as quantified in Figs. 4F and H. Therefore, it appears that UCPH-101 does not slow the recovery rate of the transient current. The most likely explanation is that, at low concentrations (<1 μM) UCPH-101 binding to EAAT1 and unbinding are very slow, with time constant of 10 s for binding, and 100s of seconds for dissociation (19). Thus, within the time course of the paired-pulse experiment (1.4 s), UCPH-101-bound and unbound transporter populations do no interconvert, and the time course of current recovery, thus, reflects that of the UCPH-101-free state. Rapid UCPH-101 application to EAAT1 in the absence of glutamate did not induce any charge movement (Fig. 4G), as expected due to the slow time course of binding.

To further increase the time resolution, allowing the separation of Na^+^ binding and translocation steps, we applied glutamate using photolysis of caged glutamate under forward transport conditions, and in the absence of permeable anions. In the absence of UCPH-101, the glutamate-induced transient current showed three phases, a rapid rising phase, and two decaying phases (Figs. S3A; black trace). The rising phase of the transient current was fitted with a time constant of τ_1_ = 0.2 ± 0.1 ms, preceding the two-exponential current decay phases. The fast decay exhibited a time constant τ_2_ = 1.2 ± 0.1 ms, the time constant of the slow decay was τ_3_ = 24 ± 5 ms. These time constants are in agreement with literature values for the homologous glutamate transporter subtype EAAC1 (43). We next pre-incubated the same cell with 50 μM UCPH-101 (a saturating concentration at steady state) for 10 s, followed by laser pulse photolysis to rapidly release glutamate. We observe that the rapidly-decaying phase of the transient current was inhibited by about 33%, but not to the full extent (Figs. S3B,C; red trace). The time constant for the current rise, τ_1_, which in EAAC1 was previously assigned to glutamate binding (44), was virtually unaffected by UCPH-101 (Fig. S3C). This result suggests that glutamate can still bind to EAAT1 in the presence of UCPH-101, as expected for a non-competitive inhibition mechanism.

In a previous report (30), the transient current’s amplitude was reduced when decreasing the external Na^+^ concentration from 140 mM to 20 mM, but the time constant of the rapidly-decaying component (time constant τ_2_) were unchanged at reduced Na^+^ concentration. This indicated that the transient signal is not generated by Na^+^ binding preceding the glutamate binding steps, but rather a conformational change associated with Na^+^ binding, most likely at the Na2 site. In contrast, the slowly-decaying phase was proposed to be associated with the glutamate translocation step (43). In agreement with this assignment of current components, the slowly-decaying phase was significantly reduced in amplitude in the presence of UCPH-101 (Figs. S3B, D; blue trace). This result suggests that UCPH-101 blocks glutamate translocation, but not other, earlier steps, such as glutamate binding and Na^+^ binding to the glutamate-bound transporter.

### UCPH-101 affects Na^+^ binding electrogenicity, but not Na^+^ affinity of the glutamate-free transporter

Does UCPH-101 binding affect how cations interact with the transporter? This question was raised from the cryoEM structures of EAAT1 in the presence of UCPH-101 (11). The structure shows Na^+^ bound only at the Na2 position (11). However, two more Na^+^ binding sites exist, Na1 and Na3 (11,15,45-49). To investigate this issue, we used voltage jumps to analyze cation binding to the *apo* (glutamate-free) transporter in the UCPH-101 bound state. As reported previously, Na^+^ binding to a glutamate-free form of EAAT2 and 3 is electrogenic (30,43,50). Step changes of the membrane potential from −100mV to +60mV resulted in transient currents (S4A (top)), which decayed with a time constant in the 0.5 ms range (140 mM Na^+^), in the absence of glutamate. Nonspecific currents were subtracted by applying 200 μM TBOA. In the presence of UCPH-101, the charge movement was reduced, but not abolished (Fig. S3B). The charge movement decreased at all voltages, but the midpoint potential was not significantly changed (Fig. S4C). These results are consistent with the interpretation that Na^+^ binding (most likely to the Na1 site) occurs with similar affinity, but with reduced electrogenicity compared to the UCPH-101-free transporter. Since Na^+^ binding presumably to the Na1 site is required for glutamate binding to occur, this result is not surprising, as the substrate can still bind in the presence of UCPH-101.

### UCPH-101 is a partial inhibitor of ASCT2

In the next paragraphs, we describe electrophysiological measurements on ASCT2 in the presence of UCPH-101, to test whether the inhibition mechanism is conserved between the EAAT1 and ASCT2 members of the SLC1 family. To verify the residues directly contributing to the UCPH-101 bound state in EAAT1 suggested by docking calculations, we used molecular dynamics (MD) simulations, to investigate the stability of UCPH-101 in its EAAT1 binding pocket (Methods). From the published EAAT1 structure in the UCPH-101 bound state (5LLM), we generated a model of EAAT1 inserted into a lipid-bilayer in a water box. After equilibration, UCPH-101 remained stable in the original binding pocket in six 100 ns simulations (Fig. 5B). We selected two conserved residues (Y127, M271; Fig. 5A and Fig. S1) to estimate distance changes to bound UCPH-101 together with RMSD calculations. Interestingly, the trajectories show that the position of UCPH-101 is stable relative to Y127 and M271 in 100 ns simulations, and these two residues stay in close contact with UCPH-101 pyran ring oxygen and the amino group nitrogen (Fig. 5D). It should be noted that UCPH-101 has two stereo-centers. For the simulations, we used the stereo-configuration analogous to the one from the EAAT1 UCPH-101-bound structure.

**Fig. 5:**
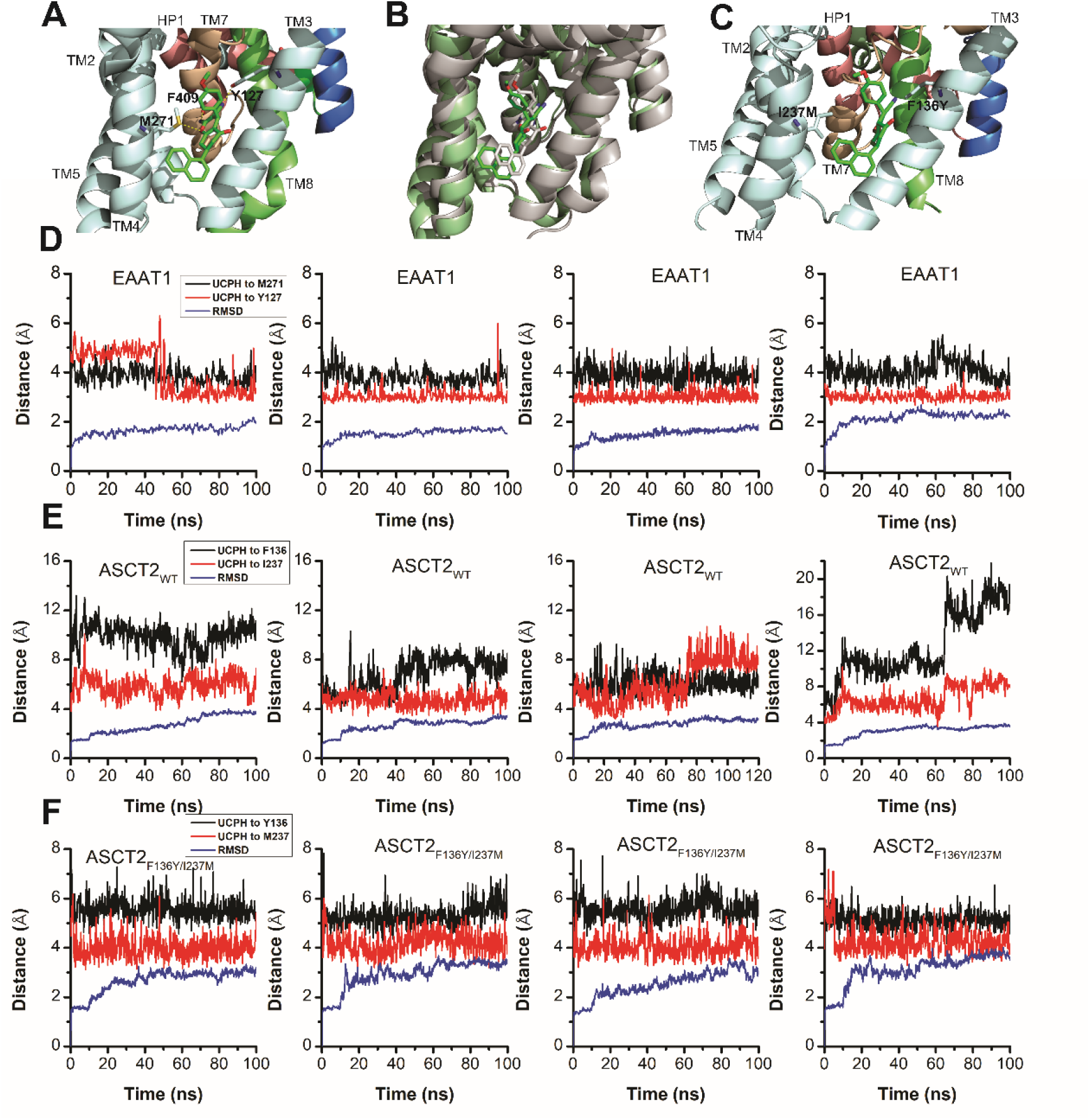
EAAT1, ASCT2_WT_ and ASCT2_F136Y/I237M_ residues contribute to UCPH-101 stability in the binding site. **(A)** Original state of the EAAT1-UCPH-101 complex from structure 5LLM (11). Y127 and M271 (EAAT1 sequence number) contribute to the EAAT1-UCPH-101 interaction. The binding state after 100 ns MD simulation is shown in **(B). (C)** Original state of the docked ASCT2_WT_-UCPH-101 complex (7BCS) with modeled side chains. Trajectories for four independent simulations runs for EAAT1, ASCT2_WT_ and ASCT2_F136Y**/I237M**_ (one UCPH-101 molecule at each subunit interface) were analyzed and are shown from **(D)** to **(F)**,(distance calculation (red, black) and RMSD distance (blue)). For distance calculations, in EAAT1, we selected atoms Y127(CA) and M271(CA) for EAAT1 and (O) and (N) for UCPH-101 (*Materials and Methods*) as reference atoms. In ASCT2_WT,_ we selected atoms F136(CZ) and I237(CD) for ASCT2_WT_ and (N1) and (O2) for UCPH-101. In ASCT2_F136Y/I237M_ distance calculation, we selected atoms Y136(OH) and M237(SD) from the transporter, and (N1) and (O2) from UCPH-101 as reference.

To investigate the stability of inhibitors in the predicted ASCT2 allosteric binding pocket, we also performed MD simulations on UCPH-101-bound ASCT2_WT_ and ASCT2_F136Y/I237M_. Residues F136Y and I237M (Fig. 5C) were selected to evaluate distance changes to UCPH-101 during equilibration. The distance of UCPH-101 is relative stable with respect to Y136 and M237 in 100 ns simulation trajectories (Fig. 5E), while a slight increase in the RMSD indicates relaxation of the structure compared to the initial, docked starting state. Consistent with our results from electrophysiology (see below) and molecular docking (Fig. 2), UCPH-101 appeared to be more stable in the ASCT2_F136Y/I237M_ binding site (Fig. 5F), where, in contrast to ASCT2 wild-type, no dissociation events were observed within 100 ns. In addition, the distance between selected atoms of UCPH-101 and ASCT2 were maintained within 6Å in the double-mutant transporter, whereas in the wild type these distances were 8-11Å (Fig. 5E), indicating that these interactions are less favorable in the wild-type transporter.

To test the predictions from docking calculations and MD simulations, we next experimentally investigated the interaction of UCPH-101 with ASCT2_WT_, and transporters with mutations to the analogous positions in ASCT2, using electrophysiological characterization of anion current, which is activated during substrate exchange. Anion conductance is an indirect measure of ASCT2 transport activity, but was previously shown to correlate with transport activity (direct amino acid transport measurements are shown below) (51). Typical anion currents, using the highly permeant anion SCN^-^, were inwardly-directed due to SCN^-^ outflow through the ASCT2 anion conductance (Fig. 6A). As expected, the anion currents increased with increasing serine concentration, with an apparent *K*_m_ of 280 ± 40 μM. When serine was co-applied together with UCPH-101, the current was inhibited, but not to 100%, even at saturating [UCPH-101] (Fig. 6B). In addition, inhibition required much higher UCPH-101 concentrations than in EAAT1. The UCPH-101 dose response relationship, at a constant serine concentration of 100 μM (Fig. 6C) demonstrates maximum inhibition of about 50% at 500 μM UCPH-101, with an apparent *K*_i_ value of 100 ± 30 μM (Figs. 6C and D). We next tested the inhibitory effect at various serine concentrations, to determine the inhibition mechanism. The maximum inhibition level varied from 50% to 18% when increasing the [serine] from 100 μM to 1000 μM (Fig. 6D). The apparent *K*_i_ was only weakly dependent on [serine]. From these results, we presume that UCPH-101 operates as a weak, non-competitive, partial inhibitor for the wild-type ASCT2 transporter (Fig. 7D).

**Fig. 6:**
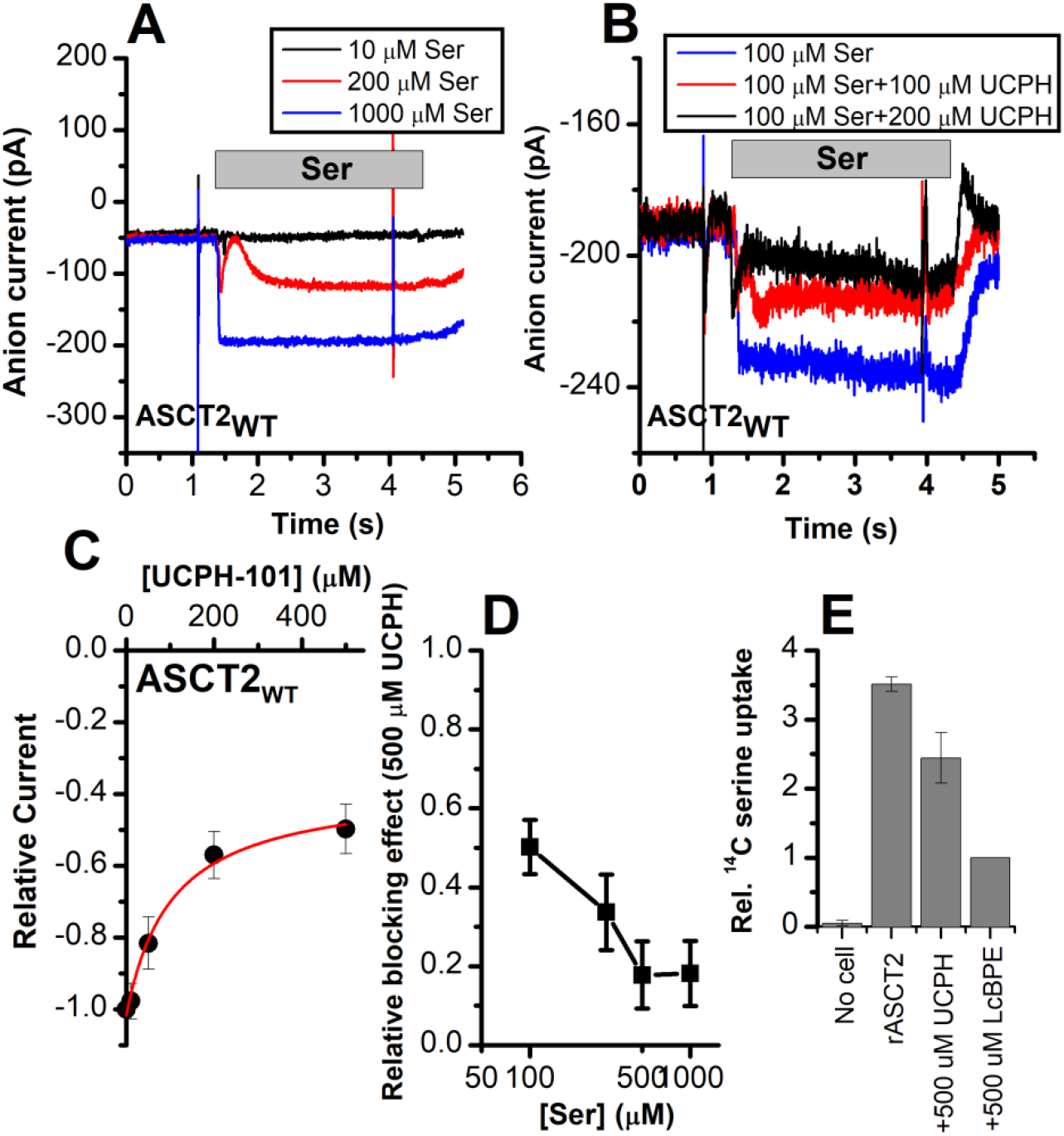
ASCT2 amino-acid induced anion current is partially inhibited by UCPH-101. **(A)** Typical serine induced ASCT2 anion current increases with increasing serine concentrations. The extracellular solution contained 140 mM NaMes, with 130 mM NaSCN and 10 mM Ser incorporated into the whole-cell recording electrode. The application time for serine is indicated by the gray bar. The apparent affinity for Ser were calculated as *K*_m_ = 280 ± 40 μM. **(B)** Same experiment as in (A) at 100 μM Ser, but in the presence of UCPH-101 at three concentrations. **(C)** UCPH-101 dose-response curve at 100 μM Ser (n=12). **(D)** Blocking effect at saturating [UCPH-101] are shown at varying Ser concentrations, the blocking effect between 100 μM Ser and 1000 μM Ser is statistically significantly different, *p*=3.14e-7. **(E)** Uptake of ^14^C serine in rASCT2-expressing HEK293 cells in the absence and presence of 500 μM UCPH-101. Competitive ASCT2 inhibitor L-cis-BPE was used at a saturating concentration to determine specific uptake by ASCT2.

**Fig. 7:**
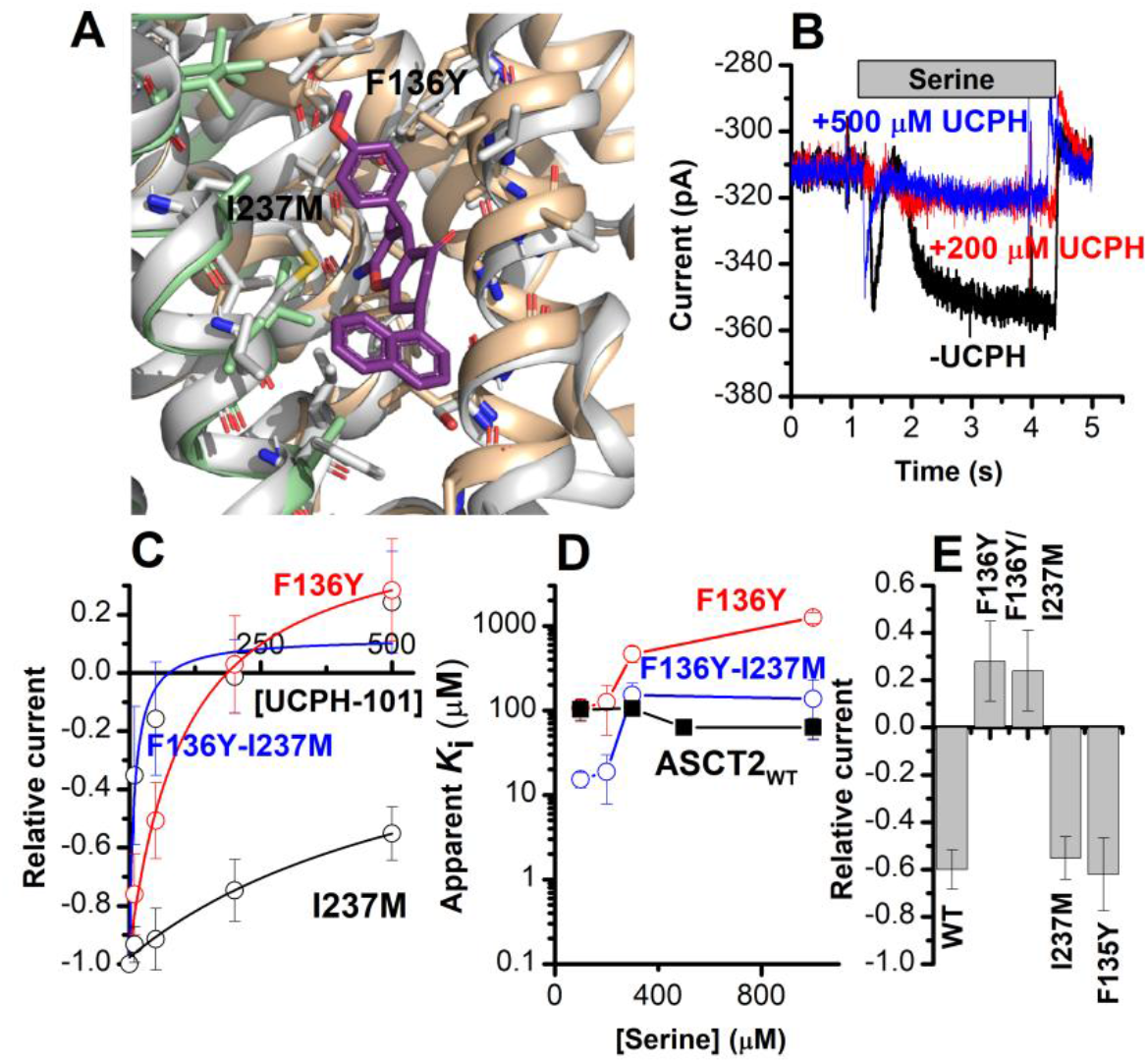
The F136Y/I237M ASCT2 double mutation restores complete inhibition of anion current by UCPH-101. **(A)** Predicted binding pose of UCPH-101 to the F136Y/I237M-mutant ASCT2 transporter. **(B)** Typical whole-cell current recordings for ASCT2_F136Y/I237M_, and UCPH-101 dose-response curves at 100μM serine. **(C)** For F136Y (red), I237M (black), and F136Y/I237M (blue) mutant transporters. Blocking effects were at a serine concentration of 100 μM and at 0 mV transmembrane potential. **(D)** UCPH-101 apparent inhibition constant (*K*_i_) plotted as a function of the serine concentration. **(E)** Comparison of steady-state current test at 100 μM Ser and 500 μM UCPH-101, wild-type ASCT2 currents without UCPH-101 were set as reference to 1, Error bars represent ± SD.

As a direct measure of amino acid transport, we next tested the effect of UCPH-101 on amino acid (L-serine) uptake. The competitive inhibitor L-*cis*-BPE was used to determine the specific component of L-serine uptake (24). We observed that serine uptake was also inhibited by UCPH-101 at a saturating concentration (Fig. 6E); however, similar to the electrophysiological data, this inhibition was not complete, with 36% of the specific uptake being blocked by UCPH-101.

To experimentally test the hypothesis, predicted by docking (see above), that the residues F136 and I237 contribute to UCPH-101 binding, four mutant ASCT2s were generated based on EAAT1, F135Y, F136Y, I237M and the F136Y/I237M double mutant (predicted docking pose for the double mutant ASCT2 is shown in Fig. 7A). Before testing with UCPH-101, the mutant transporters were functionally analyzed by whole-cell current recording experiments, to check whether the mutations altered transporter functionality. All four mutant transporters exhibited wild type-like anion currents upon substrate application. Mutant transporters with F136Y and F136Y/I237M substitutions changed the substrate selectivity, i.e. in contrast to ASCT2_WT_, anion current could not be observed after application of glutamine. In addition, the F136Y mutation decreased the serine apparent affinity from a *K*_m_ of 280 ± 40 μM (ASCT2_WT_) to 790 ± 130 μM (Table 1). None of the other mutant transporters showed a significant effect on the *K*_m_ for Ser/Ala current activation.

**Table 1.** substrate selectivity of all ASCT2 mutants

| Substrate binding affinity<br>(μM) | Ala | Ser | Gln |
| --- | --- | --- | --- |
| WT | 290±50 | 280±40 | 190±30 |
| F135Y | 300±60 | 210±50 | 190±50 |
| F136Y | 280±70 | 790±130 | insensitive |
| I237M | 230±20 | 300±70 | 270±50 |
| F136Y/I237M | 300±130 | 310±100 | Insensitive |

We next determined the effect of UCPH-101 on the mutant ASCT2s. Serine-induced anion currents were inhibited by UCPH-101 in all four mutant transporters (Figs. 7B-E), however, to varying degrees. In contrast to competitive inhibitors, which block leak anion current in the absence of substrate, application of UCPH-101 to the cells without substrate did not induce any measurable current response (Fig. 8C). Original UCPH-101 inhibition data for the double-mutant transporter and dose response curves are shown in Figs. 7B and C. In contrast to the wild-type transporter, UCPH-101 was able to block 100% of the serine-induced inward current at a concentration of 500 μM in both F136Y and the double mutant transporter (Figs. 7B and C). However, no complete block was observed in ASCT2_I237M_, and the UCPH-101 interaction was of low affinity. (Figs. 7C and D). For the F136Y mutant transporter a high apparent *K*_i_ = 1100 ± 30 μM was obtained (Figs. 7C and D). For the double mutant F136Y/I237M transporter, the apparent UCPH-101 affinity was significantly increased over the wild-type transporter. As shown in Figs. 7B and D, in ASCT2_F136Y/I237M_, UCPH-101 fully blocked the anion currents above 100 μM UCPH, resulting in a *K*_i_ value of 9.3 ± 5.9 μM. A slight outward-directed current was observed at high UCPH-101 concentrations (Figs. 7C and E), the reason for which is not known.

**Fig. 8:**
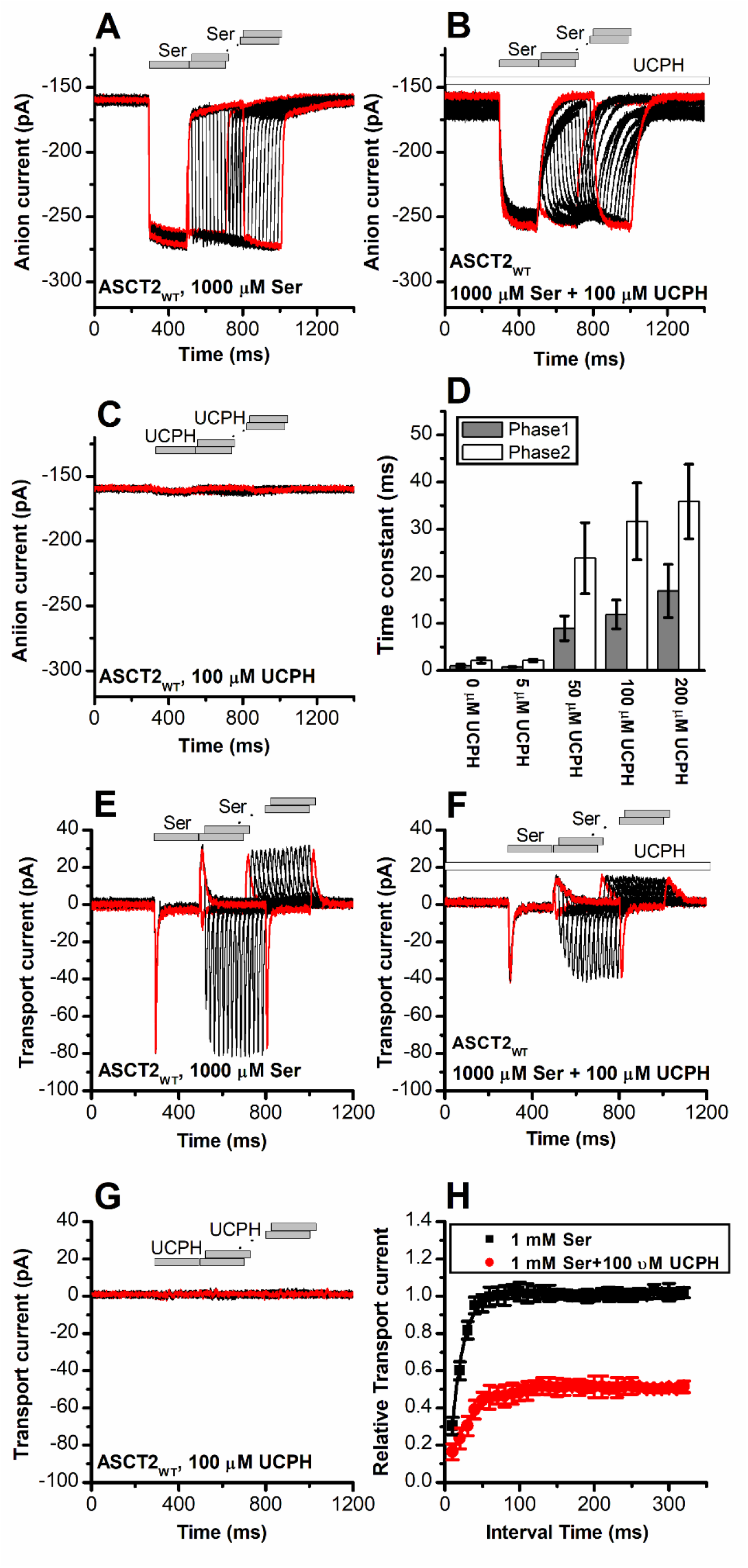
UCPH-101 slows kinetics of ASCT2 current onset and recovery after amino acid removal. **(A)** Anion currents in response to two pulses of rapid serine application (1 mM), with varying inter-pulse interval (pulse protocol shown at the top) under homo-exchange conditions. The intracellular solution contained 130 mM NaSCN/10 mM serine, the extracellular solution contained 140 mM NaMes. **(B)** Similar experiment as in (A), but in the presence of 100 μM UCPH-101 (pre-incubated for 5mins, see open bar for timing of solution exchange, top of the figure). **(C)** Application of UCPH-101 alone did not induce any currents. **(D)** Time constants for the fast and slow phase of the current recovery, for the two exponential components. **(E-G)** Experiments similar to (A-C), but for the transport component of the current (the permeant intracellular anion, SCN^-^, was replaced with the non-permeant Mes^-^ anion). **(H)** Recovery of the transient current in the presence and absence of 100 μM UCPH-101. The membrane potential was 0 mV in all experiments.

Subsequently, we compared the UCPH-101 apparent *K*_i_ as a function of the substrate concentration (Fig. 7D). For ASCT2_F136Y_, the apparent *K*_i_ increased with increasing serine concentration, which is unexpected for a purely con-competitive inhibition, but consistent with a mixed inhibition mechanism. In contrast, the apparent *K*_i_ for UCPH-101 for the F136Y/I237M double mutant transporter was largely independent of the substrate concentration, suggesting a non-competitive mechanism for block.

Finally, we tested the UCPH-101 inhibition of the double-mutant transporter at varying membrane potentials. The results are shown in Fig. S5. Serine-induced anion currents were significantly smaller than in the presence of UCPH-101 (Fig. S5B), compared to control (Fig. S5A). Inhibition was more pronounced at negative membrane potentials, as show in Fig. S5C.

### UCPH-101 slows onset and recovery of anion currents induced by substrate in wild-type and ASCT2_F136Y/I237M_

When serine was applied rapidly to ASCT2_WT_ and ASCT2_F136Y/I237M_, in the presence of intracellular SCN^-^ (substrate-induced anion current), the current was activated immediately (Figs. 8A and 9A), within the time resolution of the solution exchange (5-10 ms). In some experiments, a small, but significant transient current component was observed (Fig. 9A), due to the population of an intermediate along the translocation pathway with high anion conductance, as proposed previously (52). Unlike EAAT1 in forward transport mode, ASCT2 anion current also deactivated rapidly upon serine removal, and recovered very quickly when a second pulse of serine was applied (Fig. 8A, time of decay and recovery all within the time resolution of solution exchange). Similar results were observed for ASCT2_F136Y/I237M_ (Fig. 9A). However, when serine was applied in the presence of 100 μM UCPH-101 (pre-incubated for 5 min), steady-state anion current was inhibited, as expected, but in addition a significant slowing of anion current rise after serine application, decay after serine removal, and recovery after a second pulse of serine application was observed, for both ASCT2_WT_ (Fig. 8B) and ASCT2_F136Y/I237M_ (Fig. 9B). The time constant of the responses, which were biphasic and required fitting with two exponential components, are summarized in Figs. 8D and 9D, demonstrating a significant increase in the time constant for recovery, in particular for the fast component of the recovery. These results suggest that UCPH-101 not only inhibits current amplitude, but also slows current kinetics and serine exchange turnover (as measured by current recovery), as expected for a fast-binding inhibitor that shows rapid pre-equilibrium between the UCPH-101-bound and unbound transporter states, with respect to the substrate-induced transporter kinetics. Using pre-equilibrium conditions, the following equation can be used to describe the kinetics of the translocation equilibrium:

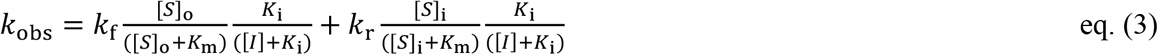

**Fig. 9:**
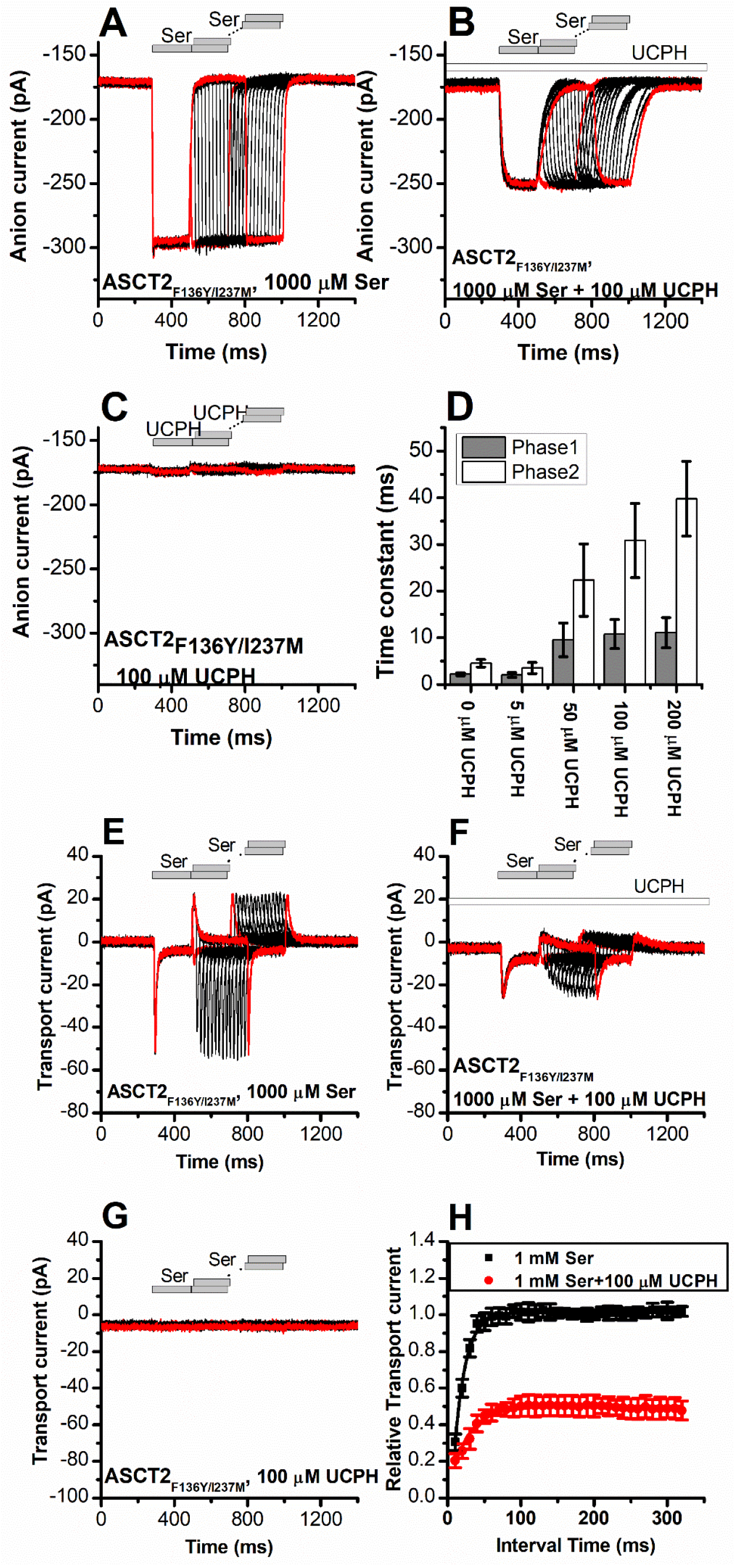
UCPH-101 slows kinetics of ASCT2_F136Y/I237M_ current onset and recovery after amino acid removal. **(A)** Anion currents in response to two pulses of rapid serine application (1 mM), with varying inter-pulse interval (pulse protocol shown at the top) under homo-exchange conditions in an ASCT2_F136Y/I237M_-expressing cell. The intracellular solution contained 130 mM NaSCN/10 mM serine, the extracellular solution contained 140 mM NaMes. **(B)** Similar experiment as in (A), but in the presence of 100 μM UCPH-101 (pre-incubated for 5mins, see open bar for timing of solution exchange, top of the figure). **(C)** Application of UCPH-101 alone did not induce any currents. **(D)** Time constants for the fast and slow phase of the current recovery, for the two exponential components. **(E-G)** Experiments similar to (A-C), but for the transport component of the current (the permeant intracellular anion, SCN^-^, was replaced with the non-permeant Mes^-^ anion). **(H)** Recovery of the transient current in the presence and absence of 100 μM UCPH-101. The membrane potential was 0 mV in all experiments.

Here, *k*_obs_ is the observed rate constant for the rise or decay of the anion current (i.e. equilibration of the translocation equilibrium), which is a sum of the rate constants for forward (*k*_f_) and reverse (*k*_r_) translocation. [S]_o_ and [S]_i_ are the external and internal substrate concentrations, respectively, [*I*] is the UCPH-101 concentration and *K*_m_ and *K*_i_ have their regular meaning. When amino acid substrate is rapidly removed, [S]_o_ becomes 0 and it is possible to isolate the reverse translocation rate constant.

### Rapid charge movements caused by ASCT2 substrate translocation are inhibited by UCPH-101

Similar to EAAT1, rapid external serine application to ASCT2 elicits transient inward transport currents (in the absence of permeable anion), which decay within 20 ms back to baseline (Fig. 8E) in the absence of UCPH-101. In contrast to EAAT1, ASCT2 is an amino acid exchanger, therefore, the experiments were performed in the exchange mode (saturating [Na^+^] and [amino acid] on the intracellular side of the membrane). Because ASCT2 amino acid exchange is electroneutral at equilibrium (equal net rate of forward and reverse charge movement), no steady state current was observed under these conditions (Fig. 8E). However, upon rapid removal of extracellular amino acid, transient outward current was observed (Fig. 8E), due to the re-equilibration of the homo-exchange translocation equilibrium to favor the outward-facing conformation in the absence of external amino acid. In contrast to the EAATs, Na^+^ apparent binding affinity to the *apo*-form of the ASCT2 transporter is high (*K*_m_ < 1 mM, (52)). Therefore, the Na^+^ binding equilibria of the amino acid-free transporter are fully saturated at 140 mM external [Na^+^], and the amino acid-induce charge movement can only be caused by Na^+^ binding to the amino acid-bound transporter, substrate translocation, or, most likely, both of those processes. Alanine and serine exchange was previously reported to be rapid in ASCT2 (25). Consistent with this proposal, transient transport current recovered much faster than in EAAT1 after amino acid removal (Fig. 8E and H), with a time constant of 15 ± 0.2 ms (Fig. 8H).

When the same paired-pulse serine solution exchange protocol was applied to ASCT2 in the presence of UCPH-101, similar transient currents were observed (inward upon serine application and outward upon removal), but the amplitude of the transient currents was significantly reduced (Fig. 8F). However, similar to the observations from the anion currents, UCPH-101 was unable to completely block transient transport current. Residual inward transient current amplitude at 100μM [UCPH-101] was 50% compared to control. As expected, UCPH-101 application alone did not generate any transient currents (Fig. 8G). Recovery time of the current was increased two-fold to 30 ± 0.4 ms at 100 μM UCPH-101. Overall, these data indicate that UCPH-101 reduces, but does not eliminate charge movements induced by Na^+^ binding to the amino acid-bound transporter, and/or substrate translocation across the membrane, in other words, the steps needed for amino acid transport by ASCT2.

Similar results were obtained for ASCT2_F136Y/I237M_ (Fig. 9). As in the case for anion current, UCPH-101 also significantly slowed the kinetics of decay and recovery of the transient transport current (Fig. 9 E-H), indicating that UCPH-101, due to decreased affinity compared to EAAT1, also acts like a rapidly-equilibrating inhibitor in the double mutant ASCT2 transporter.

### UCPH-101 effect on Na^+^ binding to the *apo*-form of ASCT2_F136Y/I237M_

It was previously shown that voltage jumps to substrate free ASCT2 result in transient charge movement from Na^+^ binding/dissociation (52) similar to EAATs (50). When a step-change in the membrane potential was applied to ASCT2_F136Y/I237M_ expressing cells (Fig. S6A, top), transient currents were observed. In these recordings, non-specific currents were subtracted by γ-(4-biphenylmethyl)-L-proline, which is commercially available, competitive inhibitor of ASCT2 (22). In the presence of UCPH-101, the transient currents were reduced at all membrane potentials (Fig. S6B). Q-V relationships are shown in Fig. S6C, suggesting that Na^+^ binding and release steps in *apo*-transporter were inhibited by UCPH-101, but not eliminated, similar to our results obtained with EAAT1 shown above (Fig. S4). Since under substrate-free conditions the translocation is not going to proceed, these results suggest that UCPH-101 likely affects Na^+^ binding to the proposed Na3/Na1 binding sites.

### Identification of a non-UCPH-101-like allosteric inhibitor of ASCT2

Our next goal was to confirm the location of the allosteric binding site by identifying compounds chemically different from UCPH-101 that interact with the ASCT2 allosteric binding site. We therefore employed virtual screening of the ZINC20 lead like library (3.8 million compounds) to identify potential binders (Methods). We analyzed the top scoring compounds for closer inspection and selected compounds that made predicted hydrogen bonding interactions to sidechains of the allosteric binding site. From the best compounds found in the virtual screening, 11 were tested experimentally. We tested the inhibitory effect at 100 µм of serine? One of these compounds (#302, Fig. 10A, docking pose shown in Fig. 10B) inhibited ASCT2 anion current (Fig. 10G, H), with an apparent *K*_i_ of 238 ± 58 µM, as well as transient transport current elicited by the transported substrate L-serine. In addition, 300 μM compound #302 inhibited the peak amplitude of ASCT2 transient transport current after rapid application of 1 mM serine to 48 ± 6 % of the control current (Figs. 10C and D). In contrast to UCPH-101, however, the presence of compound #302 did not have significant effect on the decay kinetics of the transient current, or the recovery of transient transport current after removal of amino acid substrate (τ = 19± 2.0 ms in the absence and 39 ± 1.4 ms in the presence of #302, recovery kinetics shown in Fig. 10F). As for UCPH-101, application of compound #302 alone (in the absence of serine) did not induce any current response (Fig. 10E). Notably, while compound #302 is not an optimized allosteric inhibitor of ASCT2, it provides a further validation for the location of the binding site as well as a starting point for developing future tool compounds targeting ASCT2 allosterically.

**Fig. 10:**
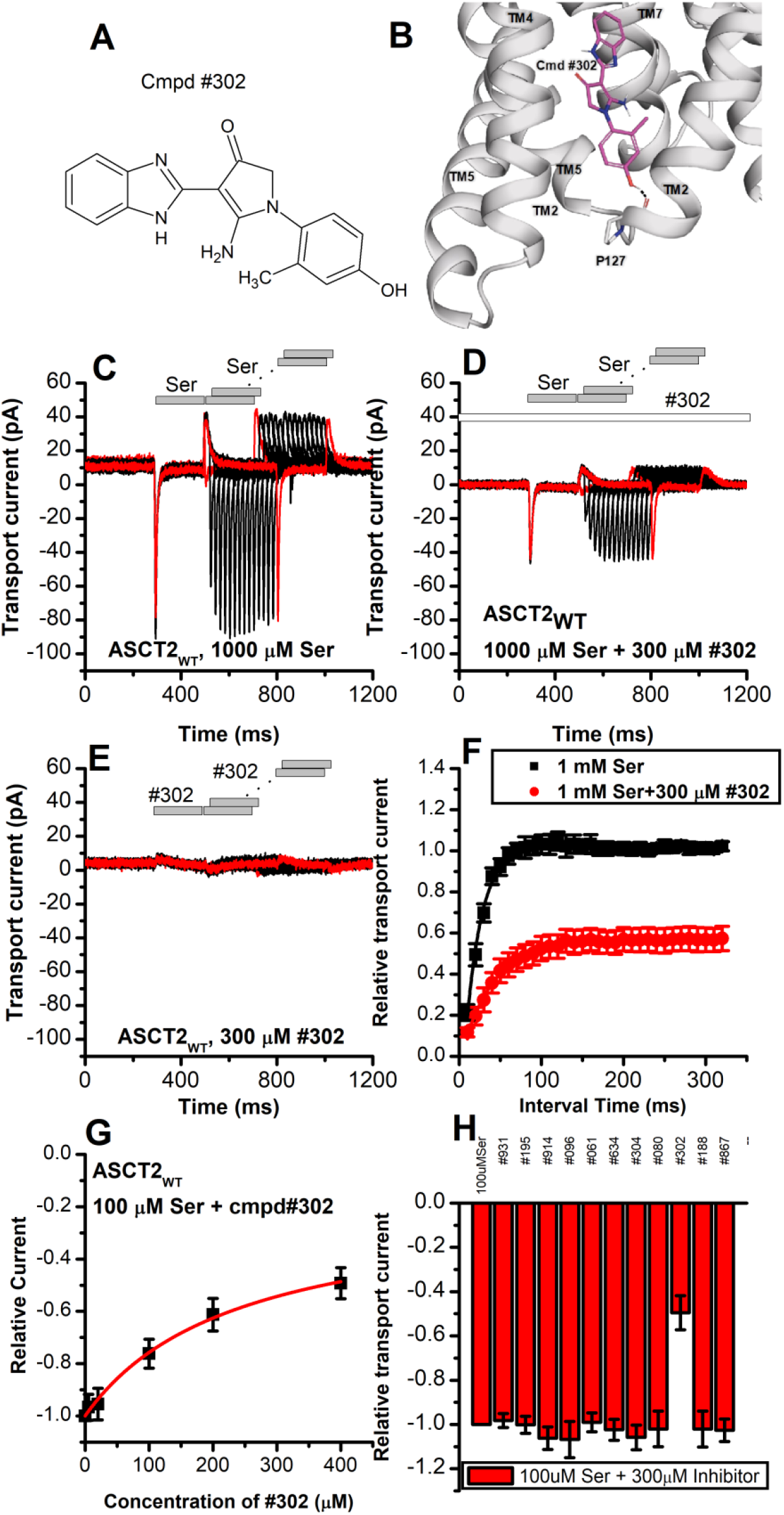
Novel Cmpd #302, identified by docking analysis, is a partial inhibitor of ASCT2 transient transport current. **(A)** Chemical structure of compound #302. **(B)** Predicted docking pose of compound #302 in the ASCT2 allosteric binding site. **(C)** Anion currents in response to two pulses of rapid serine application (1 mM), with varying inter-pulse interval (pulse protocol shown at the top) under homo-exchange conditions in an ASCT2-expressing cell. The intracellular solution contained 130 mM NaMes/10 mM serine, the extracellular solution contained 140 mM NaMes. **(D)** Similar experiment as in (A), but in the presence of 300 μM #302 (pre-incubated for 5 min, see open bar for timing of solution exchange, top of the figure). **(E)** Application of #302 alone did not induce any currents in ASCT2. **(F)** Recovery of the transient current in the presence and absence of 300 μM #302. The membrane potential was 0 mV in all experiments. **(G)** UCPH-101 dose-response curve at 100 μM Ser (n=18). **(H)** Comparison of steady-state current at 100 μM Ser and 300 μM inhibitors, wild-type ASCT2 currents without inhibitor were set as reference to 1, Error bars represent ± SD.

## Discussion

The work presented here allows us to draw four major conclusions about the mechanism of interaction of UCPH-101, and possibly other allosteric modulators, with the SLC1 family of transporters. First, UCPH-101 inhibits substrate transport by interfering with conformational changes associated with the substrate translocation reaction, rather than substrate and Na^+^ binding. Second, the general location of the allosteric binding site of UCPH-101 is conserved between the SLC1 family members EAAT1 (glutamate transporter) and ASCT2 (neutral amino acid transporter), while the transporter sidechains contributing to the binding pocket are not fully conserved. Third, occupancy of the allosteric binding site does not necessarily result in full inhibition of substrate transport, it can also lead to a partially active state, as seen for ASCT2_WT_’s interaction with UCPH-101. Our results are therefore in agreement with previous mutagenesis studies (19) suggesting that allosteric compounds may be found that modulate transport activity, rather than completely blocking it, even raising the possibility of allosteric modulators that increase transport activity by increasing the substrate translocation rate. Fourth, the allosteric ASCT2 binding site can be targeted by compounds that are non-UCPH-101-like in structure. Overall, these are novel findings that would warrant the further investigation of the allosteric binding site and the development of future, more potent compounds targeting it.

### UCPH-101 interaction with EAAT1

This report provides additional mechanistic information on the action of the allosteric inhibitor, UCPH-101 on EAATs. In agreement with previous studies, UCPH-101 shows the hallmark of a non-competitive inhibitor of EAAT1, i.e. a lack of effect of substrate concentration on the inhibitor *K*_i_. In the electrophysiology aspects, we verified that UCPH-101 affects Na^+^ interaction with the *apo*-transporters (in the absence of substrate). While it seems unlikely that Na^+^ affinity is affected, the magnitude of the voltage-induced Na^+^ binding transient currents was reduced, pointing to a reduction in the electrogenicity of the binding process. We also concluded that UCPH-101 does not affect the substrate binding steps, mainly by analysis of rapid kinetic experiments using laser-photolysis of caged glutamate. In essence, the large inward transient transport current in response to rapid glutamate application in the absence of permeant anions is only slightly affected by the presence of UCPH-101 (Fig. S3A and B), at concentrations that fully block the steady-state response. This transient current component was previously assigned to substrate-induced Na^+^ binding to the amino acid-bound transporter, and/or conformational changes associated with this binding. In order for this current component to be observed, substrate first has to bind to the transporters, which is clearly the case in the presence of UCPH-101. These conclusions are in agreement with the EAAT1 structure in complex with both UCPH-101 and a competitive inhibitor, TFB-TBOA (17), which demonstrated that the allosteric and substrate binding sites can be occupied at the same time.

In contrast to substrate binding, we propose that the substrate translocation step is the major partial reaction that is blocked in the UCPH-101-bound state. This interpretation is based on the significant reduction of the slow phase of the biphasic transient current decay in response to glutamate application, which for EAAC1 was previously assigned to the glutamate translocation step. From a structural viewpoint, this interpretation is logical, and was proposed by Canul-Tec et al. in 2017 from the EAAT1 structure in complex with UCPH-101 (11). Since binding occurs at the interface between transport and the trimerization domain, it is likely that transport domain movements relative to the trimerization domain are affected by the occupied allosteric site, either fully inhibiting those movements, or slowing them to attain partial activity.

The effect of UCPH-101 on the glutamate-induced anion current of EAAT1 fits with this interpretation. It was shown previously that UCPH-101 exhibits very slow binding and dissociation kinetics (on the 10s second time scale), much slower than the partial reactions of the glutamate-induced transport process and the turnover rate (in the 10-30 s^-1^ range). Therefore, it is likely that the UCPH-101-free and bound states interconvert slowly on the time scale of transport. Thus, UCPH-101 reduces the amplitude of glutamate-induced anion and transport current, without a significant effect on the observed kinetics associated with these reactions.

### Conserved modulatory allosteric mechanism in ASCT2

The development of novel inhibitors are essential steps for potential treatment of diseases, especially for ASCT2 transporter. In previous reports, overexpression of ASCT2 was observed in several cancer cell lines (7-10). Among the several published ASCT2 inhibitors, most of them target the substrate binding site. However, allosteric modulators would be useful because of the independence of their inhibitory effect on substrate concentration. The known allosteric binding pocket of EAAT1 as well as the biomedical importance of ASCT2 prompted us to investigate whether a similar allosteric mechanism is present in ASCT2; and, indeed, UCPH-101 was found to interact with this binding site. Therefore, the importance of this report lies in the generation of a new structural model proposed here to develop future novel allosteric modulators with the potential for higher affinity and specificity for ASCT2 in future compounds.

In the EAAT members of the SLC1 family, UCPH-101 was found to be specific for EAAT1. For example, in wild-type Glt-1 (EAAT2), no inhibition was observed, unless EAAT1/EAAT2 chimeras were generated that contained the EAAT1 N-terminal six transmembrane domains (19). In contrast to the EAATs, however, it was not known whether UCPH-101 can interact with the ASCT members of the SLC1 family. Our results reported here are, to our knowledge, the first to show that UCPH-101 inhibits ASCT2, providing evidence for the existence of the allosteric binding site in the same location as in EAAT1. While UCPH-101 is only a partial inhibitor of wild-type ASCT2, and its apparent inhibition constant is relatively high (low affinity, in the range of 100 μM), it demonstrates the principle of allosteric inhibition, or modulation of ASCT function through the allosteric binding site. Importantly, we were able to restore complete inhibition of substrate-induced anion current in mutant ASCT2 transporters, in which the proposed UCPH-101 binding site was modified to resemble that of EAAT1. In the double mutant ASCT2_F136Y/I237M_ the affinity was also increased compared to the wild-type transporter, although not reaching the high affinity levels seen in EAAT1.

As expected, UCPH-101 shows the hallmarks of a non-competitive inhibitor, except for the F136Y mutant transporter, in which a mixed inhibition mechanism was observed. The reason for this is not known, but somehow the UCPH-101 binding site in this mutant transporter must communicate with the substrate binding site. A proposed kinetic mechanism for inhibition of EAAT1/ASCT2 by UCPH-101 is shown in Fig. 11, in analogy to previously published alternating access mechanisms for the SLC1 family (53,54). Here, UCPH-101 can bind to all of the states the transporter can access during the transport cycle, including substrate and Na^+^-bound states. The reaction that is affected by occupation of the allosteric site is the translocation reaction, or reactions closely associated with it, such as Na^+^ binding to the substrate-bound transporter. In EAAT1, due to the high affinity of the compound, equilibration of the UCPH-101-bound and unbound states is slow, thus, UCPH-101 inhibits the transient and steady-state currents, with little effect on apparent transport kinetics. In essence, UCPH-101 acts by removing active EAAT1 transporters from the pool, thus inhibiting the overall transport rate. On the other hand, equilibration of the UCPH-101-bound and unbound states is fast in ASCT2, therefore, kinetics of substrate-induced current activation and recovery are slowed in the presence of inhibitor. Thus, the kinetic mechanism shown in Fig. 11 can account for the subtleties of kinetic differences in UPCPH-101 effects between EAAT1 and ASCT2.

**Fig. 11:**
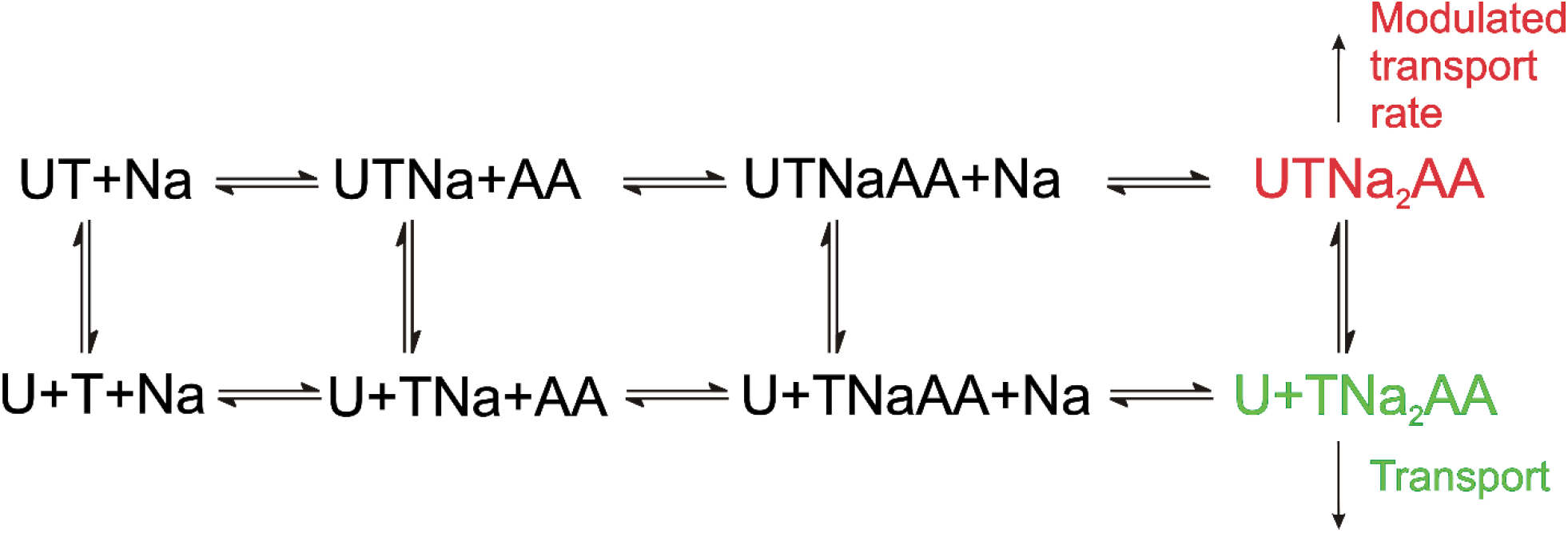
Proposed simplified mechanism of inhibitors (U) targeting the allosteric binding site interacting with ASCT2 and EAAT1 transporters (T). The amino acid substrate is abbreviated as AA. Potential charges of the cation (Na^+^) and the transporter/substrate were omitted for simplicity, and only two of the three Na^+^ binding steps are shown. For UCPH-101 interaction with EAAT1 the UCPH binding/dissociation process (transition from the first to the second row of the scheme) is slow, while this transition is fast for ASCT2, accounting for the differential effect of UCPH-101 on transport kinetics of both transporters.

One limitation with current compounds targeting ASCT2 for cancer therapy is that they are amino-acid like compounds targeting the substrate binding site, and thus have poor pharmacokinetics. Interestingly, we were able to identify a compound with a structure unrelated to UCPH-101, compound #302, which inhibited wild-type ASCT2, although at relatively high concentrations and, like UCPH-101, showing only partial inhibition of the transient transport current elicited by substrate application. This compound was found from virtual screening against the putative allosteric binding site. Like UCPH-101, compound #302 is relatively hydrophobic, and harbors an amino function. However, it also has a OH group, suggesting that additional hydrophilic interaction may be important in the allosteric binding site. These results further support the presence of the allosteric binding site in ASCT2 and suggest that it may be possible to identify allosteric inhibitors of ASCT2 with novel scaffolds, opening new avenues for future structure activity relationship (SAR) studies, to improve affinity and specificity.

ASCT2 has recently received attention due to its involvement in neutral amino acid homeostasis in cells, in particular cancer cells. It was shown that down-regulation of ASCT2 function either by siRNA, or by competitive inhibitors, can block the growth of cancer cells in cell-line models, as well as in vivo tumor transplants (55,56). Competitive inhibitors have the disadvantage of becoming less effective as the amino acid concentration rises. Having allosteric inhibitors available would result in amino acid concentration independent block of ASCT2 function, which would be much more desirable for pharmacological applications. The existence of the allosteric binding site on ASCT2 raises the possibility that such compounds could be developed, which, in contrast to UCPH-101 would result in complete inhibition of transporter function.

## Conclusions

The results presented in this report reveal a conserved allosteric inhibition mechanism for SLC1 members for the first time. The results suggest that the major effect of the compound, once bound to the binding site at the interface between the transport and trimerization domain, is to block reactions associated with substrate translocation and anion conductance activation. In contrast, other partial reactions, such as substrate and Na^+^ binding can still take place, although the electrogenicity of Na^+^ binding is reduced, pointing to a slight conformational change of the substrate-free transporter state induced by UCPH-101. While UCPH-101 is a partial and weak inhibitor of wild-type ASCT2, complete inhibition could be restored by mutating the UCPH-101 binding site to one that resembles EAAT1. Therefore, our results suggest that ASCT2 also harbors the allosteric binding site, however, with differences in the amino acid side chains contributing to the site. These results raise the possibility that this allosteric binding site can be exploited by future compounds specifically targeting ASCT2. Our results involving the novel inhibitor compound #302 further confirm the location of the allosteric binding site in ASCT and suggest that compounds with varying chemical scaffolds could be found that modulate, or completely inhibit SLC1 activity. In light of the emerging role of ASCT2 as an anti-cancer drug target, the future characterization of this allosteric site appears promising.

## Acknowledgements

This study was supported by a grant from the National Institutes of Health (http://www.nih.gov.eresources.mssm.edu) (R01 GM108911) to A.S. and C.G.; and T32 CA078207 to R.A.G.; and the R15 GM135843-01 awarded to C.G.

## Suppl. Materials

**Fig. S1.**
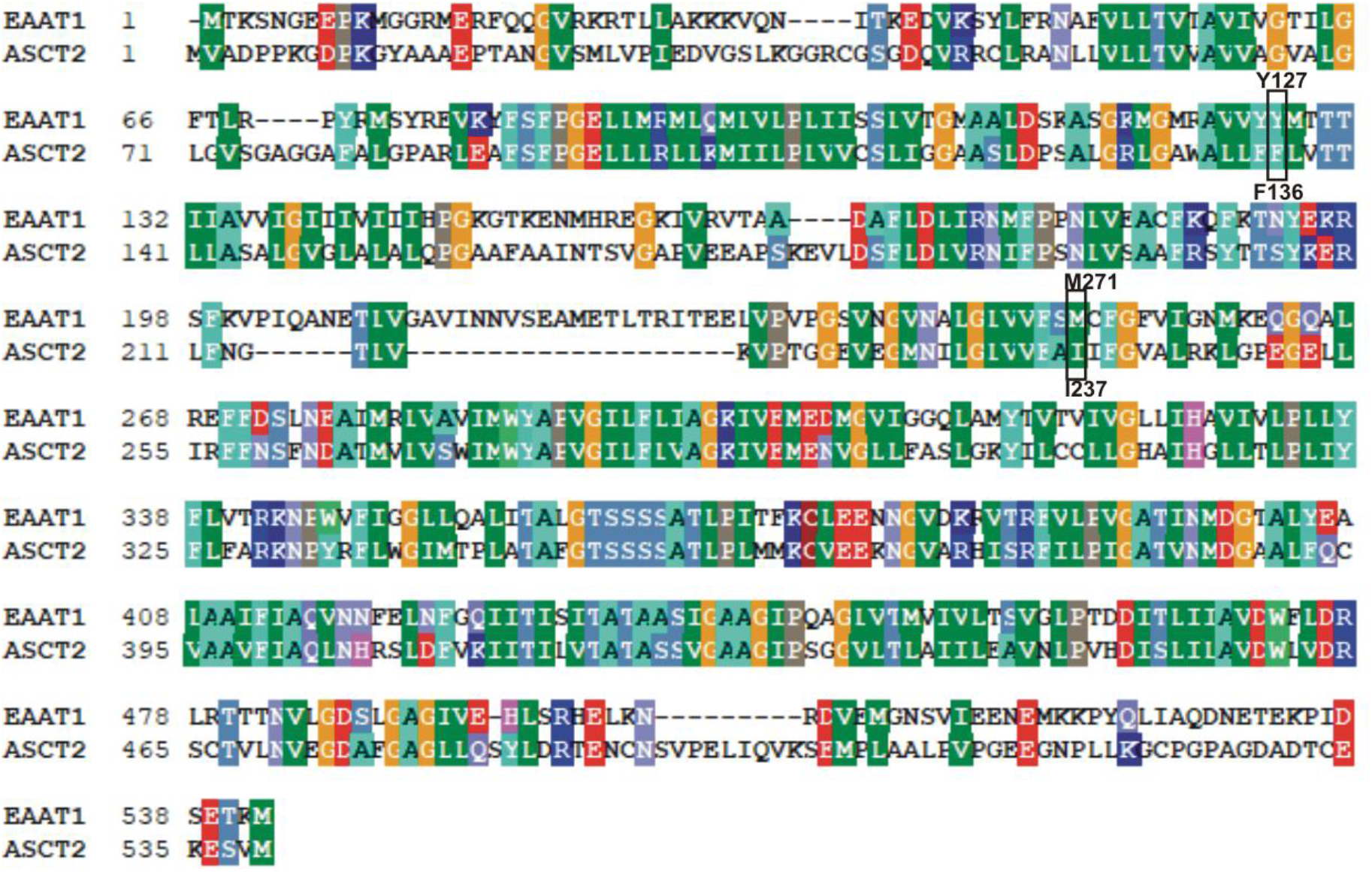
EAAT1/ASCT2 sequence alignment. Sequence alignment was perfrormed with Bioedit software, total alignment length: 571 amino acid, 220 Identical residues, 109 Similar residues. 38.53% percent identity and 57.62% percent similarity. The position of mutants F136Y, I237M are highlighted.

**Fig. S2:**
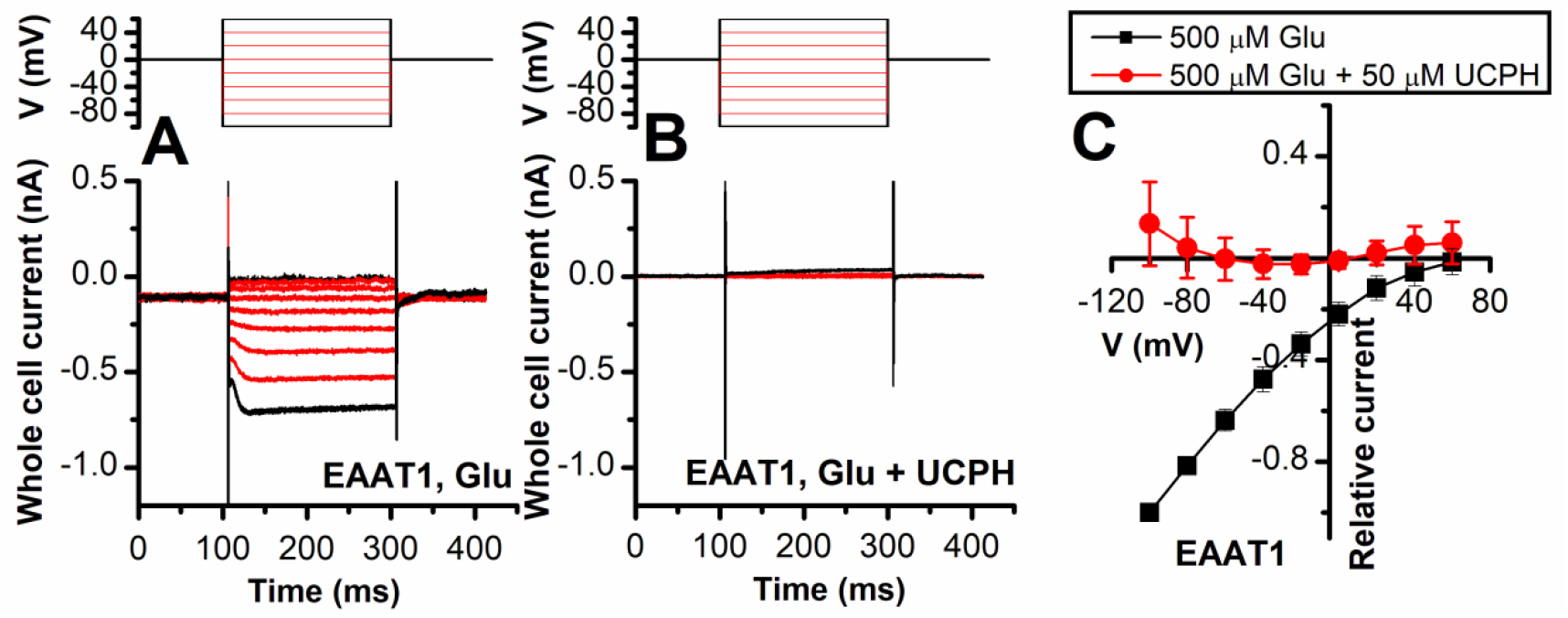
EAAT1 anion current is inhibited by UCPH-101 independent of voltage. Voltage-dependence of EAAT1 anion currents under forward transport conditions, activated using 500 μM extracellular Glu (n=13) **(A)** or 500 μM Glu in the presence of 50 μM UCPH-101 (n=14) **(B)**. The voltage-jump protocol is show in the top panels. The extracellular solution contained 140 mM NaMes, the intracellular solution contained 130 mM KSCN. The anion current-voltage relationship at steady-state is shown in **(C)**. Non-specific background currents were subtracted using 200 μM TBOA.

**Fig. S3:**
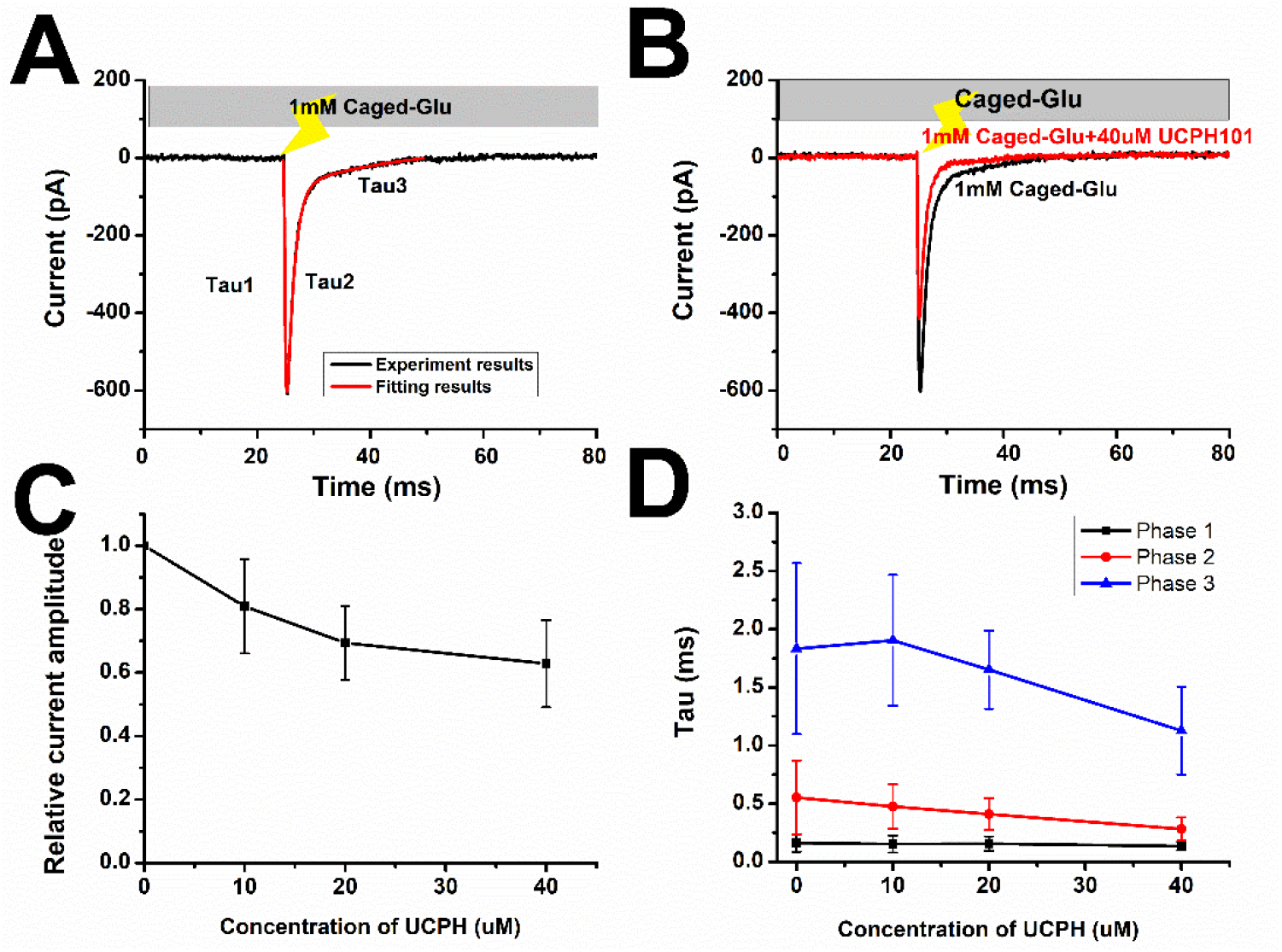
UCPH-101 inhibition blocks mainly the slow phase of the EAAT1 glutamate-induced pre-steady state current response. **(A)** Typical transient transport current signal in response to photolysis (indicated by arrow) of 1 mM caged-glutamate, Currents were fitted with a sum of three exponentials (red), τ1 = 0.2 ± 0.1 ms, τ 2 = 1.2 ± 0.1 ms, τ3 = 24 ± 5 ms). The extracellular solution contained 140 mM NaMes, the intracellular solution 130 mM KMes. **(B)** The red trace shows a similar experiment in the presence of 40 μM UCPH-101. **(C)** Relative transient current amplitude at different concentrations of UCPH-101. **(D)** Time constants of the three exponential components plotted as a function of [UCPH-101]. The components are colored in black, red and blue, respectively.

**Fig. S4.**
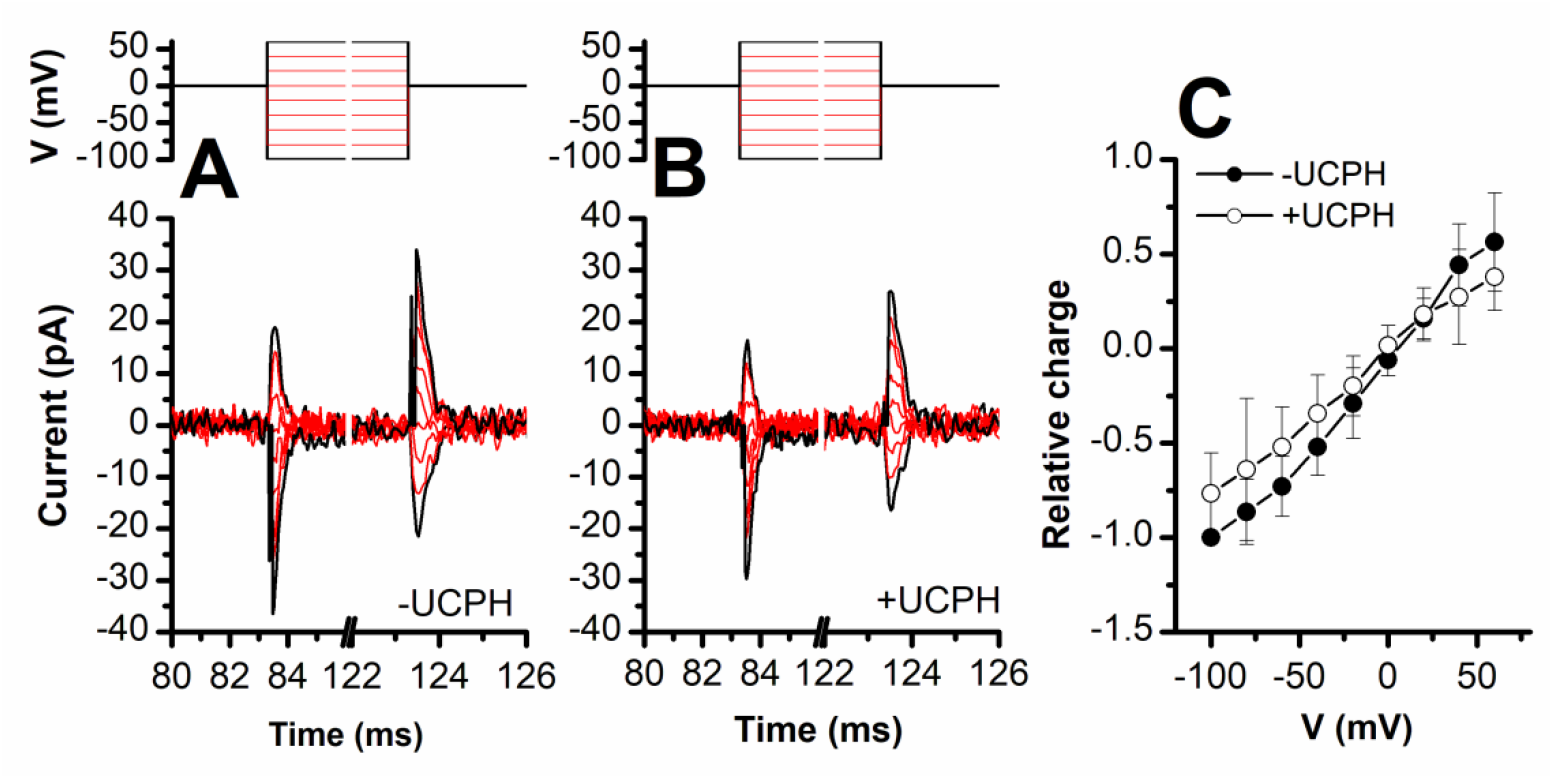
UCPH-101 does not eliminate, but reduces capacitive charge movement caused by Na^+^ binding to the *apo*-form of EAAT1. Voltage jumps (protocol shown in top panel) were used to disturb the Na^+^ binding equilibrium in the *apo*-form of EAAT1 (absence of glutamate). **(A)** and **(B)** show the transient currents caused by Na^+^ moving into and out of the binding site in the absence and presence of UCPH-101 (50 μM), respectively. Experiments were performed using extracellular solutions containing 140 mM NaMes and intracellular solutions containing 130 mM KMes. Unspecific currents were subtracted using 200 μM TBOA. **(C)** Charge movements were integrated from the transient current signals and plotted as a function of the membrane potential. The closed and open circles show relative charge movements in the absence (n=8), and presence of 50 μM UCPH-101 (n=8), respectively.

**Fig. S5:**
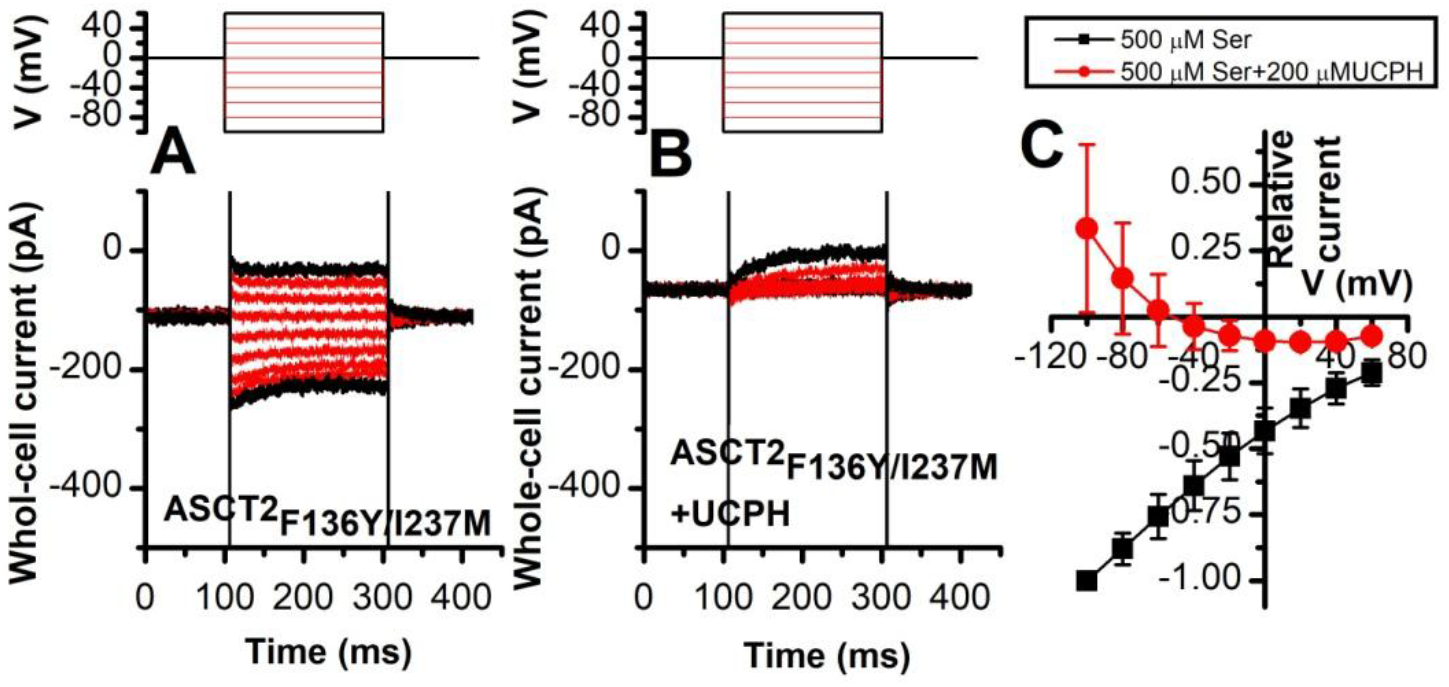
ASCT2_F136Y/I237M_ anion current is inhibited by UCPH-101 independent of voltage. Voltage-dependence of ASCT2_F136Y/I237M_ anion currents under forward transport conditions, activated using 500 μM extracellular Ser (n=13) **(A)** or 500 μM Ser in the presence of 200 μM UCPH-101 **(B)**. The voltage-jump protocol is show in the top panels. The extracellular solution contained 140 mM NaMes, the intracellular solution contained 130 mM NaSCN and 10 mM Ser. The anion current-voltage relationship at steady-state is shown in **(C)**. Background currents were obtained by application of 140mM NMG-Mes and then subtracted to generate the ASCT2_F136Y/I237M_ specific currents.

**Fig. S6.**
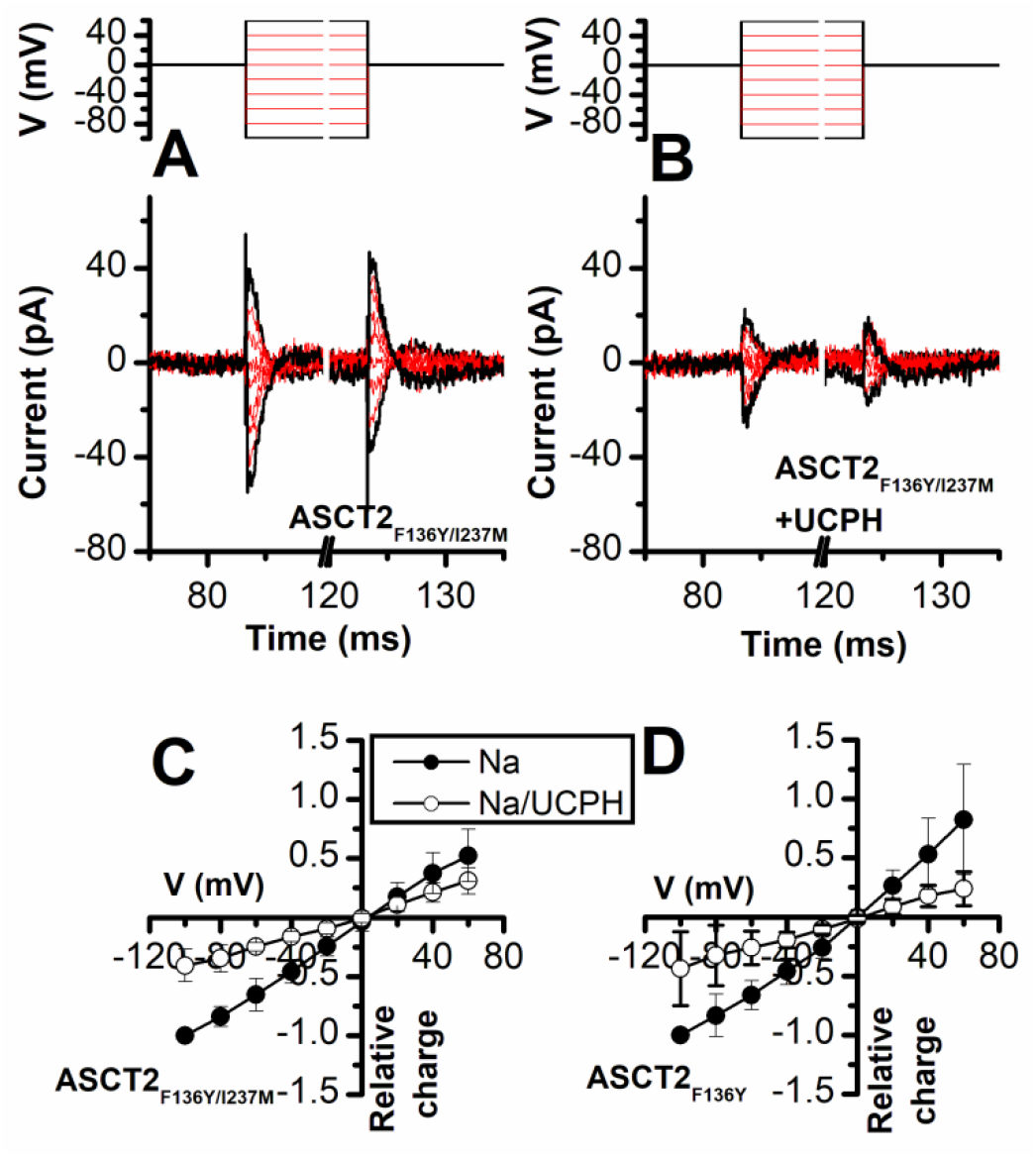
UCPH-101 does not eliminate, but reduces capacitive charge movement caused by Na^+^ binding to the *apo*-form of ASCT2_F136Y_ and ASCT2_F136Y/I237M_. Voltage jumps (protocol shown in top panel) were used to disturb the Na^+^ binding equilibrium in the *apo*-form of ASCT2_F136Y/I237M_ (absence of amino acid substrate). **(A)** and **(B)** show the transient currents caused by Na^+^ moving into and out of the binding site in the absence and presence of UCPH-101 (200 μM), respectively. Experiments were performed using extracellular solutions containing 140 mM NaMes and intracellular solutions containing 130 mM NaMes and 10 mM serine. **(C, D)** Charge movements were integrated from the transient current signals and plotted as a function of the membrane potential for ASCT2_F136Y/I237M_ (C, n = 18) and ASCT2_F136Y_ (D, n = 18) in the absence (black squares), and presence (red circles) of 200 μM UCPH-101, respectively.

## Notes

### Competing Interest Statement

The authors have declared no competing interest.

